# Large clones of pre-existing T cells drive early immunity against SARS-COV-2 and LCMV infection

**DOI:** 10.1101/2022.11.08.515436

**Authors:** Martina Milighetti, Yanchun Peng, Cedric Tan, Michal Mark, Gayathri Nageswaran, Suzanne Byrne, Tahel Ronel, Tom Peacock, Andreas Mayer, Aneesh Chandran, Joshua Rosenheim, Matthew Whelan, Xuan Yao, Guihai Liu, Suet Ling Felce, Tao Dong, Alexander J. Mentzer, Julian C. Knight, Francois Balloux, Erez Greenstein, Shlomit Reich-Zeliger, Corrina Pade, Joseph M. Gibbons, Amanda Semper, Tim Brooks, Ashley Otter, Daniel M Altmann, Rosemary J Boyton, Mala K Maini, Aine McKnight, Charlotte Manisty, Thomas A. Treibel, James C. Moon, COVIDsortium Investigators, Mahdad Noursadeghi, Benny Chain

## Abstract

We analyzed the dynamics of the earliest T cell response to SARS-COV-2. A wave of TCRs strongly but transiently expand during infection, frequently peaking the same week as the first positive PCR test. These expanding TCR CDR3s were enriched for sequences functionally annotated as SARS-COV-2 specific. Most epitopes recognized by the expanding TCRs were highly conserved between SARS-COV-2 strains, but not with circulating human coronaviruses. Many expanding CDR3s were also present at high precursor frequency in pre-pandemic TCR repertoires. A similar set of early response TCRs specific for lymphocytic choriomeningitis virus epitopes were also found at high frequency in the pre-infection naïve repertoire. High frequency naïve precursors may allow the T cell response to respond rapidly during the crucial early phases of acute viral infection.

**One-Sentence Summary:** High frequency naïve precursors underly the rapid T cell response during the crucial early phases of acute viral infection.

## Main Text

T cell responses to vaccination (*1*) and acute infection (*2*) are often fast, with the magnitude of the response peaking within one to two weeks of exposure, and substantially preceding maximum antibody responses. T cells may therefore combine with innate immunity to control microbial growth early in infection, before humoral immunity is fully developed. The mechanisms which underly this early clonal expansion remain poorly defined. In this study, we exploit a granular longitudinal sampling of peripheral blood cells which were collected during, and in some cases preceding acute infection with SARS-COV-2, during the first few weeks of the pandemic (*3*). The dynamic changes in the T cell receptor repertoire which are observed during this period reveal mechanisms that allow the speed of the T cell adaptive response to compete with that of innate immunity and viral replication during the crucial early phases of this acute viral infection.

There is an increasing interest in the role of T cells in providing protection against SARS-COV-2 infection and disease (*4*). HLA-peptide binding and in vitro peptide-driven expansion have provided a comprehensive catalogue of T cell epitopes (*5*, *6*), which span most of the viral open reading frames (ORFs). Single cell protein and transcriptomic analyses have provided complementary information on memory and effector T cell function (*7*, *8*). There have also been many attempts to link T cell responses to clinical outcome, although distinguishing cause and effect in these observational studies is challenging, especially because SARS-COV-2 disease is often associated with lymphopoenia. Nevertheless, there is some evidence that T cell responses may be associated with better clinical outcomes (*9*–*11*) and this has fueled the development of more T-cell centric vaccines, especially in individuals with impaired humoral immunity(*12*).

Despite the wealth of published studies on SARS-COV-2, however, dynamic analysis of immune responses in the early phases of infection are still rare(*13*). Indeed, detailed dynamics of the earliest responses are rare in any human infection. Experimental infection studies (e.g. to influenza) have provided some important data in this regard, but prior exposure is often a confounder in this setting (*14*). We have previously reported on the COVIDSortium (*15*), a longitudinal study of London-based health care workers, established very early in the first wave of the COVID pandemic, which provides a detailed granular view of the peripheral blood compartment during the first few weeks of infection (*3*). We have used this sample set to demonstrate a remarkably fast CD4 and CD8 T cell response following SARS-COV-2 infection, which parallels the innate interferon response, and substantially precedes humoral immunity (*16*). In the present study, we investigate in more detail this early T cell response, using robust quantitative T cell repertoire sequencing to chart the evolution of the anti-viral response at the level of the clonotype.

Global TCRseq provides an unbiased survey of the T cell response, which is complementary to more antigen-specific methodologies which depend on predefined targets such as peptides or HLA multimers. We identify a wave of early T cells, which are enriched for sequences independently annotated as SARS-COV-2-specific. Many of the CDR3 sequences are found in the pre-pandemic repertoires of healthy individuals, and we leverage this information to develop a statistical framework to infer that they are present at high clonal frequency in pre-pandemic repertoires. Remarkably, we extend these observations to similar early expanding epitope-specific T cell receptors (TCRs) in LCMV infection, and demonstrate that many of these sequences can be found at high abundance in naïve repertoires prior to and independently of exposure to lymphocytic choriomeningitis virus (LCMV). The ability to rapidly recruit high abundance TCRs commonly found in the naïve TCR repertoire may be a common strategy to cope with the response to novel microbial pathogens.

## Results

We sequenced the T cell repertoire of whole blood RNA samples collected at different time points (see below) from 41 health care workers (HCW) who tested PCR+ for SARS-COV-2 infection, and 6 HCW who remained PCR and seronegative throughout the study (sample collection summarized in Fig S2). The samples were collected at weeks 0 - 4 (acute) and week 12-14 (convalescent). The median number of samples collected for each individual was 4, ranging from 1 to 6. The majority of infections registered a PCR+ test at the first sample, but in 12 individuals, samples predating the first PCR+ sample were obtained. The first 3-4 weeks of sample collection coincided with the first wave of the SARS-COV-2 pandemic in London, and no additional cases of PCR+ confirmed infection or seroconversion were observed in the subsequent 2-3 months of collection. We sequenced a total of 14.6 million TCR alpha and TCR beta genes, with a median 73,000 TCR alpha and 94,000 TCR beta sequences per sample (Fig S3A). The number of alpha and beta sequences per sample were highly correlated (Spearman correlation = 0.93, Fig S3B), providing additional confidence in the quantitative robustness of the pipeline.

We first measured the richness (defined as number of distinct sequences, divided by total number of sequences in sample) and Shannon diversity (which incorporates a measure of the frequency distribution of the sequences) of the repertoires at different time points in both PCR+ and PCR- individuals. For PCR+ individuals we plotted the values relative to the week at which the individual first became PCR+ (Fig S4). However, there were no detectable effects of infection on the global parameters of the TCR repertoire.

We therefore focused on identifying individual TCRs which changed significantly in abundance within the first 5 weeks of the study (see Materials and Methods and (*15*) for further details). In some pairwise comparisons a clear population of expanded TCRs was observed (Fig 1A). In others, especially where the first sample available was already PCR+ , a population of contracting TCRs was observed (Fig S5A). In these cases, we made the assumption that we had missed the expansion phase, but observed the contraction phase of the initial T cell response. We therefore combined all up and downregulated TCRs, which we refer to as expanded TCRs, and removed MAIT cells (defined as TRAV1-2 paired with TRAJ12, TRAJ20 or TRAJ33) and iNKT cells (identified as TRAV10 paired with TRAJ18 ) from further analysis, leaving 4075 expanded TCR alpha sequences and 5458 expanded TCR beta sequences. In control uninfected individuals there were few expanding or contracting TCRs (Fig S5B). The richness and diversity of the set of expanded TCRs increased at the time of first PCR+ in infected individuals (p-value < 0.05 for all time points compared to pre-infection, Wilcoxon signed-rank test), but showed no consistent dynamics in uninfected individuals (Fig 1B). Note that the time axis was rescaled relative to the week at which that individual became PCR+ (renamed as week 0).

**Fig 1.**
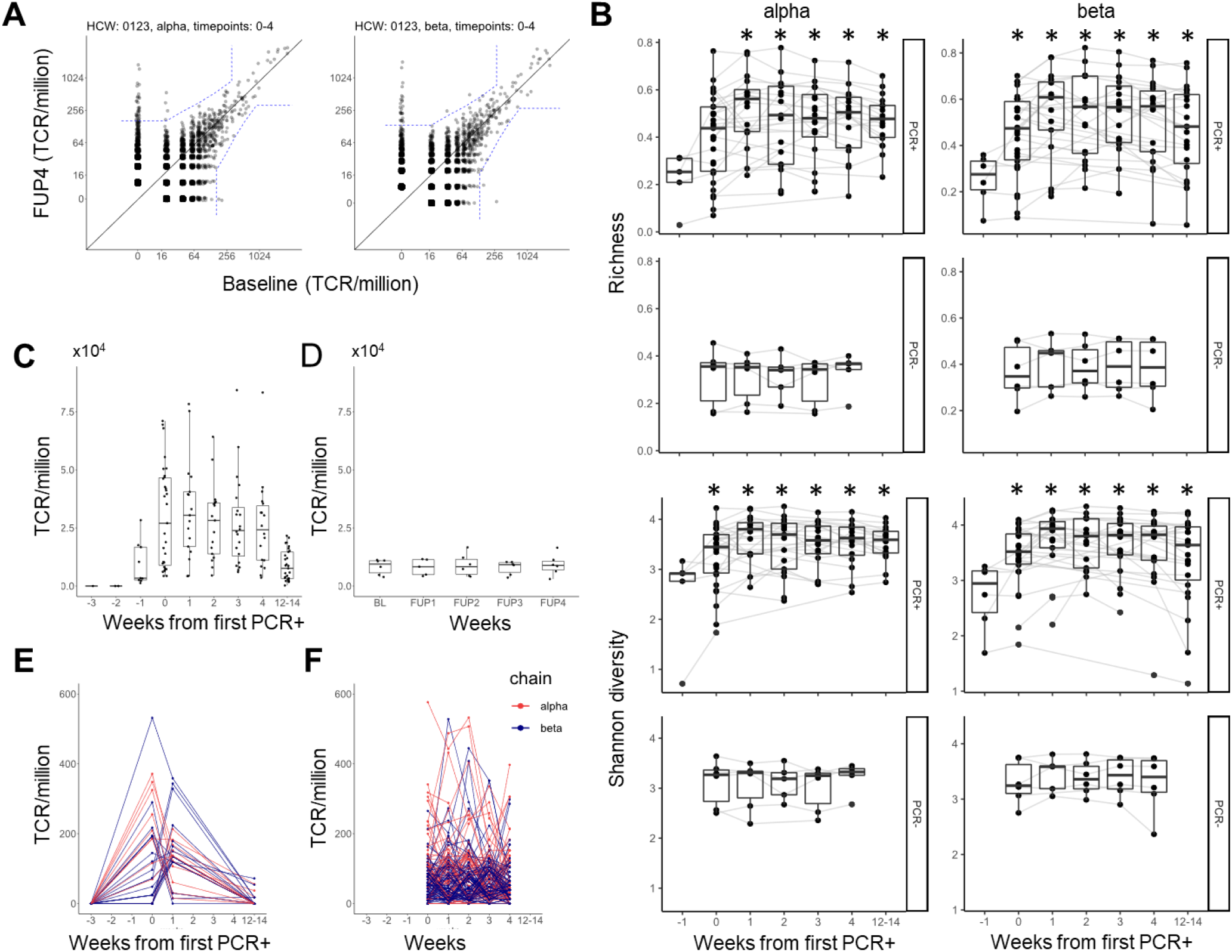
An early transient wave of TCR expansion associated with infection with S2. A. An example of a pairwise comparison between two time points, showing TCRs expanding between baseline and follow up week 4 (FUP4) repertoires. The individual (ID 123) became PCR+ at follow up week 1 (FUP1). Each point is an individual TCR sequence, plotting its abundance at FUP4 versus its abundance at Baseline. All abundances are normalized to number of TCRs per million. The dashed blue line indicates the significance threshold calculated as described in M&M. All TCRs which fall outside the dashed line are considered as expanded (contracted). B. The richness and Shannon diversity of the set of expanded (contracted) TCRs at each time point, subsampled to similar dataset size. For PCR+ individuals, the x-axis is rescaled relative to the week at which they first became PCR+ (this is week 0). For PCR- individuals the weeks correspond to baseline and subsequent follow-ups at weeks 1-4. * p-value < 0.05, Wilcox test compared to week -1. C. The sum of the expanded TCRs at each timepoint, for each infected individual. The timepoints have been recalibrated relative to the week of the first PCR+ test (this is defined as week 0). The boxplots show median, interquartile (box) and 95% (whiskers) range. D. As for C, but for controls who did not become PCR+. E. The dynamics of individual TCRs (TCRalpha in red, TCRbeta in blue) in one individual (ID 101). F. As for E but for a PCR- (ID 17). The dynamics for each individual HCW are shown in supplementary data.

We plotted the combined abundances of all expanded TCRs for each individual (an estimate of total clonal expansion) as a function of time (Fig 1C). The magnitude of the response varied considerably between individuals, but showed a common dynamic. The response was already strong at week 0 and fell by week 14. In some individuals responses could be detected even in the week prior to becoming PCR+. The magnitude of the response of the “expanded” TCRs in the PCR- seronegative controls showed no evidence of any common dynamic pattern (Fig 1D). For comparison we show the dynamics of the anti-spike (Fig S6A) and anti-nuclear protein (Fig S6B) antibody titres, which rose more gradually and slowly over the first 14 weeks, as described in detail previously (*34*).

An example of the time course of an individual HCW, for whom we had samples pre as well as post infection, shows more clearly the rapid expansion of individual TCR alpha and TCR beta sequences, which were mostly not detected pre-infection, because of sampling only a very small proportion of the repertoire, but maximal at either week 0 (week first PCR+) or week 1 (Fig 1E). Although the magnitude of the T cell response returns towards baseline by week 14, the expanded TCRs in this individual remain significantly above their starting abundance (Fig S7). For comparison, the timecourse of TCRs in an individual who did not become PCR+ show no clear pattern over the weeks of sampling (Fig 1F). The individual time courses for TCRs for all HCW are shown in Fig S8.

In summary, analysis of the dynamics of the TCR repertoire over the first 4 weeks identified a set of TCRs whose abundance increased rapidly and transitorily around or shortly after the time of SARS-COV-2 detection.

### TCRs expanding early are enriched for sequences associated with SARS-COV-2 recognition

We collected a set of 7694 (7632 unique across all sets, Fig 2A) functionally annotated TCRs, combining from a public TCR database VDJDb (*35*), a published study (*36*) and a set of TCRs we identified by tetramer sorting or in vitro peptide expansions (*23*). The functional affinity of the TCRs varied over a broad range, but included several high affinity clones (Fig S9). 239 (141 unique) CDR3s from the expanded TCR set matched with CDR3 sequences in the annotated TCR sequences (3.1%, Fig 2B). The majority of matches were with alpha chains. Same size random sets of non-expanded TCRs from the HCW repertoires matched an average of 4.2 (0.06%) annotated TCRs, significantly (more than 10 fold) less that the number of annotated sequences in the expanded set (P<0.0001, Fisher’s exact test). There were only 7 matches of SARS-COV-2 annotated CDR3 with the set of expanded TCRs from the control (PCR-) repertoires.

**Fig 2.**
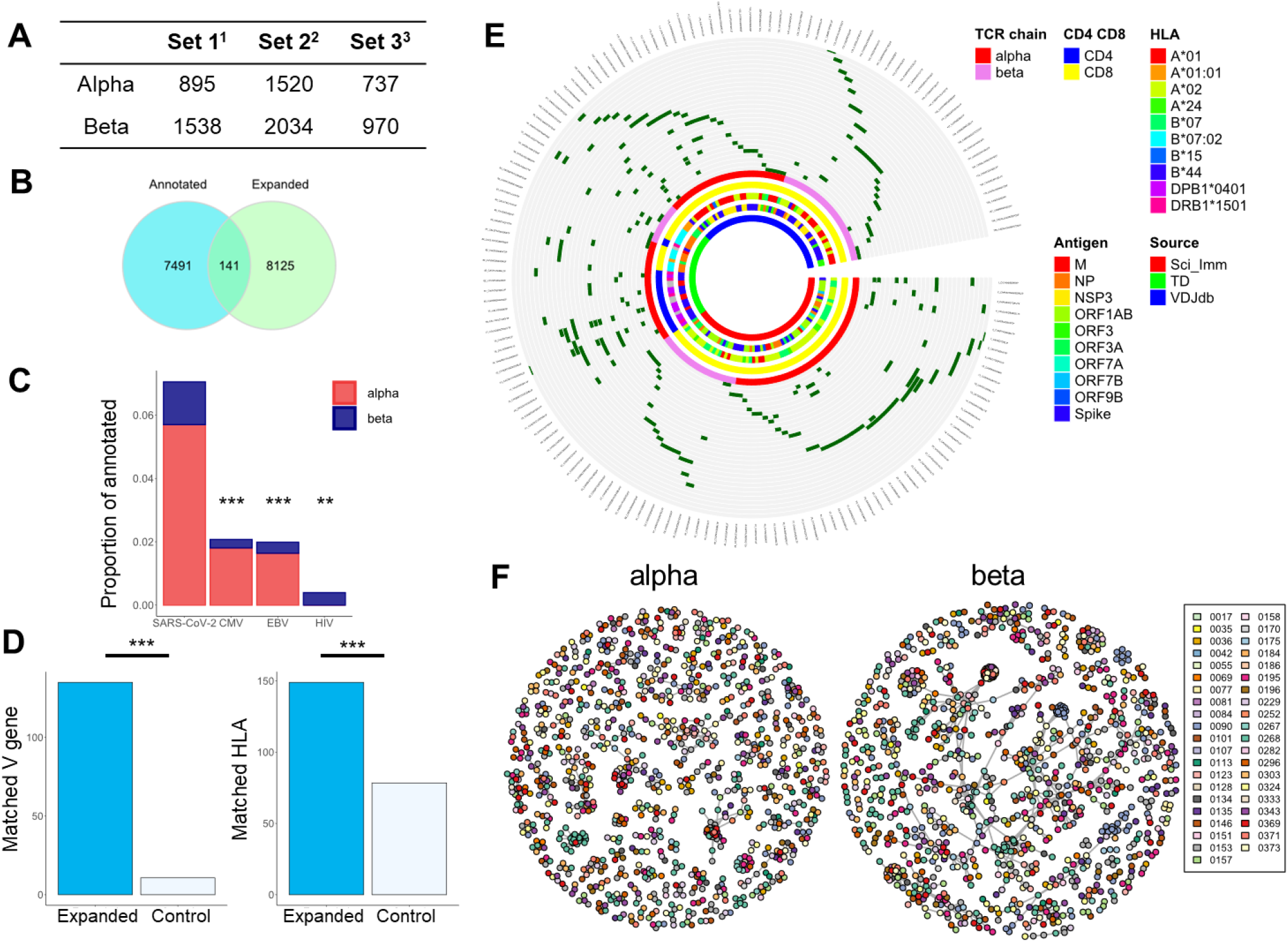
Functional annotation for SARS-COV-2 and sequence similarity characterise the early wave of expanded TCRs in PCR+ individuals. A. We collected a set of TCR sequences which had been functionally annotated for recognition of SARS-COV-2 epitopes. The set was compiled from three data sets (3423) (*36*) (3423). The set combines paired alpha/beta sequences and unpaired sequences. The Table shows the number of alpha and beta sequences in each set. B. The intersection between the expanded TCRs as defined in Fig 1 (blue), and the SARS-COV-2 annotated TCRs as in panel A (green). C. The proportion of annotated TCRs for each of four viruses which are observed in the expanded set of TCRs defined in Fig 1. SARS-COV-2 annotated TCRs are found significantly more often than those for CMV, EBV and HIV (*** p <0.0001, ** p<0.001, Fisher’s exact test). D. Left panel. Blue shading - the number of expanded TCR CDR3s which share the same V gene as the respective identical annotated TCR CDR3. Empty bars – the number of matches after random shuffling of the expanded set with respect to their V gene. Right panel. Blue shading – the number of annotated TCRs whose restricting HLA gene matches at least one allele of the individual in whom the identical expanded CDR3 is detected. Empty bars – the number of matched TCRs after random shuffling of the annotated TCRs with respect to their HLA restriction (***p<0.0001, Fisher’s exact test). E. A circus plot showing distribution of the annotated expanded TCRs in the cohort, and the metadata associated with each TCR. Each segment is a unique CDR3. Each circle is a different individual. Dark green segments correspond to an annotated expanded TCR. The inner circles show which CDR3 are alpha or beta; which CDR3 are derived from CD4 or CD8 T cells; the restricting HLA allele for each CDR3; the target antigen recognised by the annotated TCR; and the source of the CDR3. F. A graph representation of the TCR alpha and beta sequence similarity. Each node is a CDR3 sequence, and edges connect all nodes with a string kernel similarity index of greater than 0.76 For TCRalpha and 0.72 TCRbeta. Each individual is shown in a different colour. Only clusters of three or more connected nodes are shown.

We examined whether the expanded sets were also enriched for CMV annotated or EBV annotated TCR, since these viruses are prevalent in the general population (Fig 2C). The enrichment for SARS-COV-2 annotated TCRs was significantly higher than for CMV or EBV. As a comparison, we looked for matches with HIV annotated TCRs, since this virus is not likely to be prevalent in the general population. The proportion of matches with the HIV set was smaller than CMV or EBV, but the lack of annotated HIV-specific TCRalpha sequences in the database complicates interpretation.

Sharing of an identical CDR3 sequence between SARS-COV-2 annotated TCRs and the expanded set of TCRs identified in this study does not necessarily guarantee that the antigen target of the CDR3s is the same. However, we observed that the expanded CDR3 in our data was associated with the same TCR V gene as reported in the annotated set (Fig 2D, left). In addition, the HLA of the individuals in whom we observed expanded matched CDR3s was associated with the HLA restriction reported for the epitope reported for each annotated TCR (Fig 2D, right). The sharing of CDR3 sequence, V gene and HLA restriction are together suggestive of shared epitope recognition.

The distribution of annotated expanded TCRs among the PCR+ HCW cohort, and the associated metadata available for each epitope are shown in Fig 2E. Almost every PCR+ individual contained at least one annotated expanded TCR, and many contained several. The annotated matched TCRs included both CD4+ and CD8+ cells, were restricted by a variety of HLA alleles and recognized epitopes from a variety of structural and non-structural viral proteins. The time course of the annotated TCRs broadly matched that of the total set of expanded TCRs, with the majority of TCRs peaking the week of the first PCR+ test (Fig S10).

A number of expanded TCRs (including some expanded annotated TCRs, Fig 2D) were found in more than one individual. Since TCRs with similar CDR3 sequence often recognize the same antigen (*37*–*39*) we looked for sharing of similar, as well as identical expanded TCRs between individuals. Similarity was measured using a triplet kernel as described previously (*40*) and in M&M. We observed that a substantial proportion of the expanded TCRs formed inter-individual clusters of similar sequence, indicating that similar TCRs were frequently expanded in different individuals following SARS-COV-2 infection (Fig 2F). The proportion of control (non-expanded) TCRs forming clusters was much less (Fig S11). Some of the clustered expanded TCR sequences were also functionally annotated, and where more than one annotated TCR was found within a cluster, the target antigen was generally the same (Fig S12).

Taken together, the data presented in Fig 2 support the hypothesis that the early wave of expanding TCRs which follow infection with SARS-COV-2 represent a set of virus specific TCRs, many of which share similar or identical sequences in different individuals.

### Expanding TCRs associated with SARS-COV-2 infection are abundant in healthy pre-pandemic repertoires

The peak in TCR expansion observed around the time of PCR+ virus detection suggests a very rapid T cell response. We explored the possibility that the rapid expansion of TCRs arises from sequences which are present at higher than average frequency in the pre-infection repertoire. We developed a statistical framework to address this hypothesis using a large set of published PBMC repertoires (Fig 3A) .

**Fig 3.**
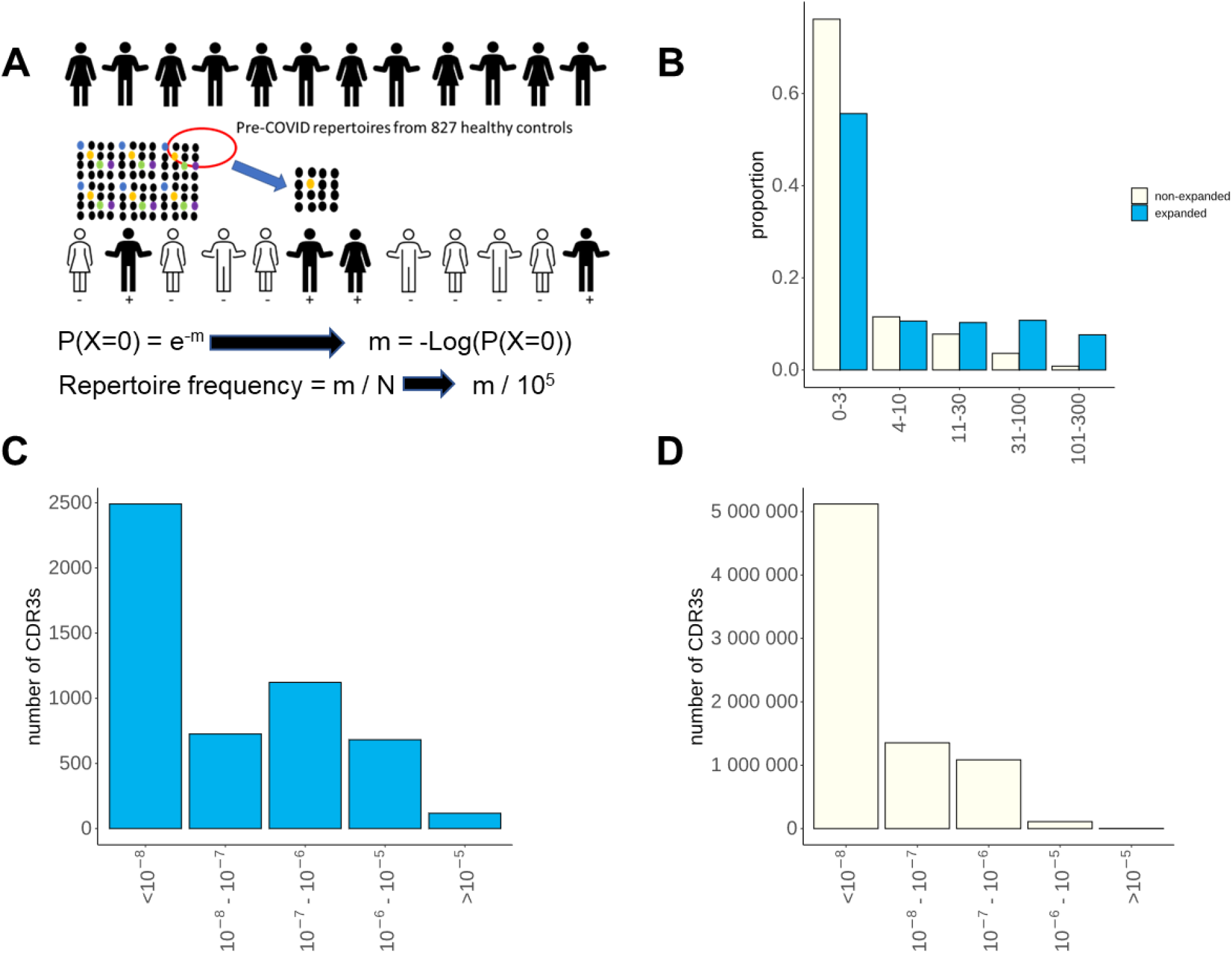
Sharing and frequency estimates of SARS-COV-2 expanded TCR . A Schematic of statistical inference of TCR frequencies from abundance data in pre-pandemic repertoires. m is the mean number of times a particular TCR is present in a given sample, given that that TCR is NOT detected in a proportion P of the 786 pre-pandemic repertoires examined. The estimated frequency is given by m/N, where N is the average number of TCRs sequenced in each sample. B. Sharing of expanded TCRbeta or non-expanded control TCRs across the 786 repertoires of the Emerson data set, collected and sequenced several years before the SARS-CoV-2 pandemic. The x axis shows the number of individuals each TCR is observed in. The y axis shows the proportion of the expanded TCRs with a given sharing level. C. The estimated frequency distribution of the 2648 expanded TCR beta sequences which are found in 2 or more individuals of the Emerson data set. The frequency of each expanded TCR was estimated using equation 2, as discussed in the text. TCR which were found less than twice in the Emerson data were assigned a frequency of <10^-6^ and are represented by the column closest to the y axis. D As for C, but for non-expanded TCRs from the SARS-COV-2 repertoires.

We determined the proportion of repertoires from a cohort of 786 individuals collected and sequenced pre-SARS-Cov-2 (referred to here as the Emerson data set (*32*), which contain each of the expanded CDR3 sequences defined in our study. Since each repertoire from the Emerson set contains only a small sample of the total repertoire in each individual (on average 1.8*10^5 sequences per sample), the inter-individual sharing must reflect a high abundancy of the shared TCR in the sample. The average abundance of the TCR in the repertoire can be estimated for all those TCRs which are seen at least twice in the Emerson data set using the Poisson probability distribution. Specifically, the proportion of the individuals which do NOT contain a particular sequence (p(X = 0)) is given by

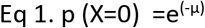

where μ is the average number of times a TCR is observed in a sample. The average number of TCRs sequenced per sample in the Emerson data is 1.8*10^5. Therefore the frequency of the TCR in the repertoire, in TCRs/million can be estimated by

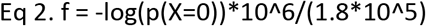

As shown in Fig 3B the proportion of these SARS-COV-2 naïve individuals in whom we found each of the SARS-COV-2 expanded TCR is much greater than for non-expanded TCRs from the same repertoires. 2648 (52%) of the 5139 unique TCRbeta expanded sequences (the Emerson data is restricted to beta sequences) were observed two or more times in the Emerson data set. The estimated frequency distribution of these expanded TCRs estimated using equation 2 is plotted in Fig 3C, and compared to non-expanded TCRs. The mean frequency of the expanded set is 1.8 per million, over a thousand times higher than 1 per 10^9, which as discussed above is the estimated frequency of individual TCR sequences in the naïve repertoire.

### Expanded TCRs in infected hosts cannot be explained by cross-reactivity to other coronaviruses

We have previously demonstrated that human TCR repertoire has a very broad range of TCR frequencies (*41*). High frequency precursor T cells could therefore arise either because they arise from naïve T cells with a higher than average frequency or due to cross-reaction (*9*, *42*). The samples were collected during the first wave of SARS-COV-2 in the UK, and the response could not therefore be attributed to pre-existing memory responses against SARS-COV-2. We therefore explored the hypothesis that the early wave of TCRs represented a restimulation of cross-reactive memory cells present as a result of exposure to common circulating human corona-viruses (hCoVs). In order to test this hypothesis, we analysed the 65 peptide epitopes which were recognized by the 141 TCRs which were functionally annotated as SARS-COV-2 specific and were also identified as early expanders in our data set (as detailed in Fig 2). The similarity between these peptide epitopes and the homologous sequences from the major strains of circulating hCoVs is shown in Fig 4A. In general, with the exception of one NSP12 (polymerase) epitope, the SARS-COV-2 sequences in the hCoVs showed substantial differences from their homologous sequences in SARS-COV-2, making it unlikely that the TCRs specific for these sequences would recognize the equivalent peptide from the circulating hCoVs. In contrast, the epitopes recognized by these early annotated TCRs showed extremely high homology between the different variants of SARS-COV-2 (Fig 4B). Within spike protein, for example, the early T cell epitopes were distinct from the mutations which are known to affect antibody recognition (Fig S13)(*43*). This analysis does not in any way preclude the existence of T cells which cross-react between SARS-COV-2 and circulating hCoVs as suggested in other studies (*9*, *44*). Rather, the analysis argues that the observed rapid expansion of SARS-COV-2-specific TCRs can occur even in the absence of any such cross-reaction.

**Fig 4.**
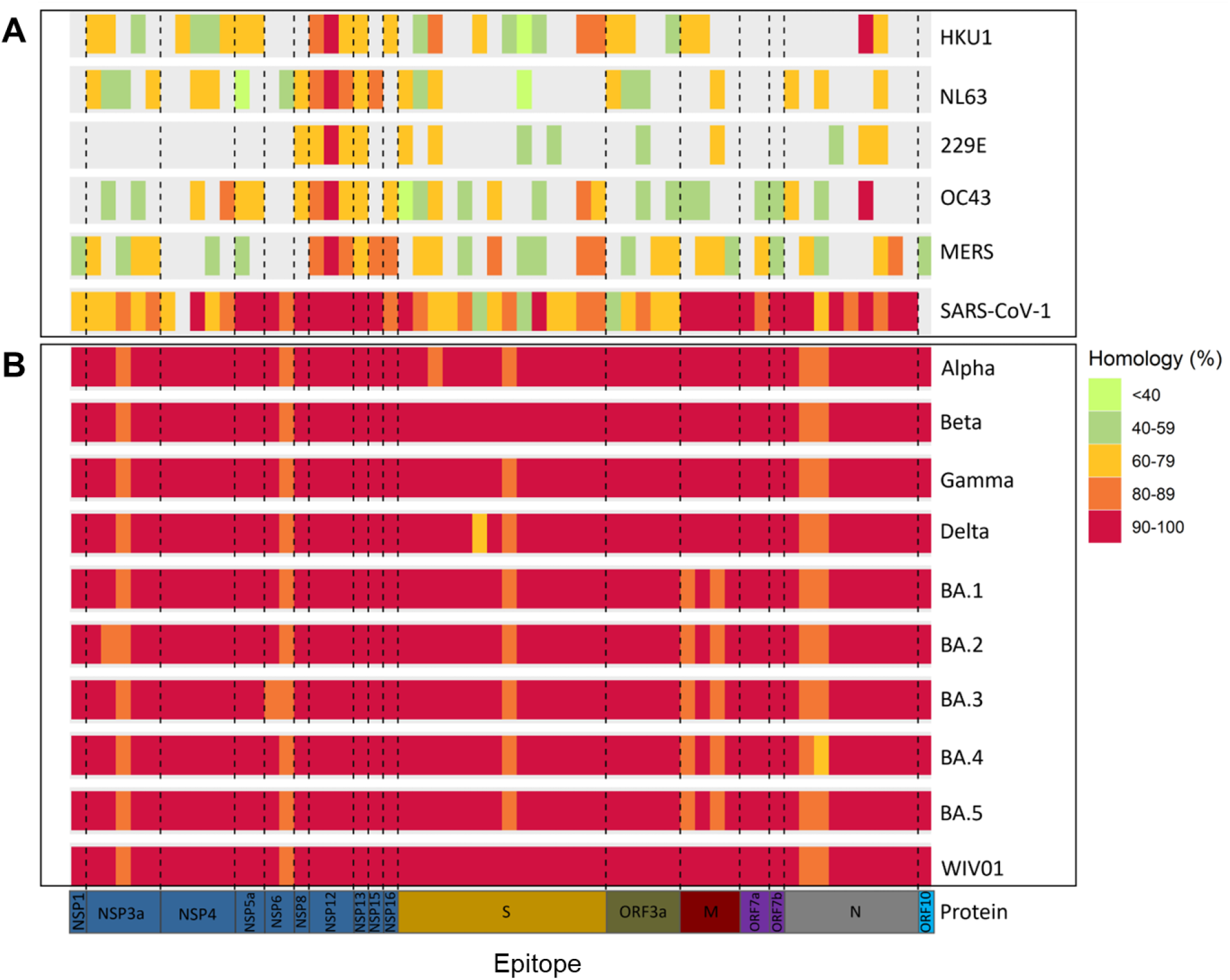
The homology between T cell epitopes of SARS-COV-2 and circulating human coronaviruses. A. Circulating coronaviruses B Different strains of SARS-COV-2 Each column is a functionally defined T cell epitope of SARS-COV-2 which is recognised by a TCR whose sequence is found among the set of TCRs expanding following infection with the virus (as defined in Fig 1). The degree of homology is shown by colour coding.

### High frequency LCMV specific TCRs in the naïve repertoire

The number of viable cells collected at early timepoints of the COVIDSortium study limited the types of analysis we could carry out, and hence the ability to fully investigate the alternative hypothesis that the expanding TCRs we identified were derived from large pre-existing T cell clones in the naïve repertoire. However, we reasoned that the phenomenon we observed for SARS-COV-2 might be common to other viral infections, and might reflect fundamental features of the naïve T cell repertoire. We therefore examined the abundances of TCRs from C57BL/6 mice exposed to LCMV. This model has been studied extensively, and several dominant and sub-dominant epitopes have been described (*45*–*47*). We harvested spleens from mice at day 8 and day 40 post infection with saline or LCMV. In contrast to the SARS-COV-2 data, we did not have to rely on dynamics to identify virus specific T cells. Instead we directly isolated antigen specific T cells by sorting with fluorescent MHC tetramers bound to four well-described LCMV epitopes GP66, GP92, NP205 and NP396. An aliquot of spleen cells from each animal was separately fractionated into naïve (CD62L+, CD44-), central memory (CD62+, CD44-) and effector (CD62-, CD44+) phenotype as described in Materials and Methods. We sequenced the TCR repertoire of each antigen-specific and non-specific sample, and then searched for antigen specific TCR (tetramer-bound) sequences in the repertoire of the bulk naïve and effector populations (Fig5 A).

**Fig 5.**
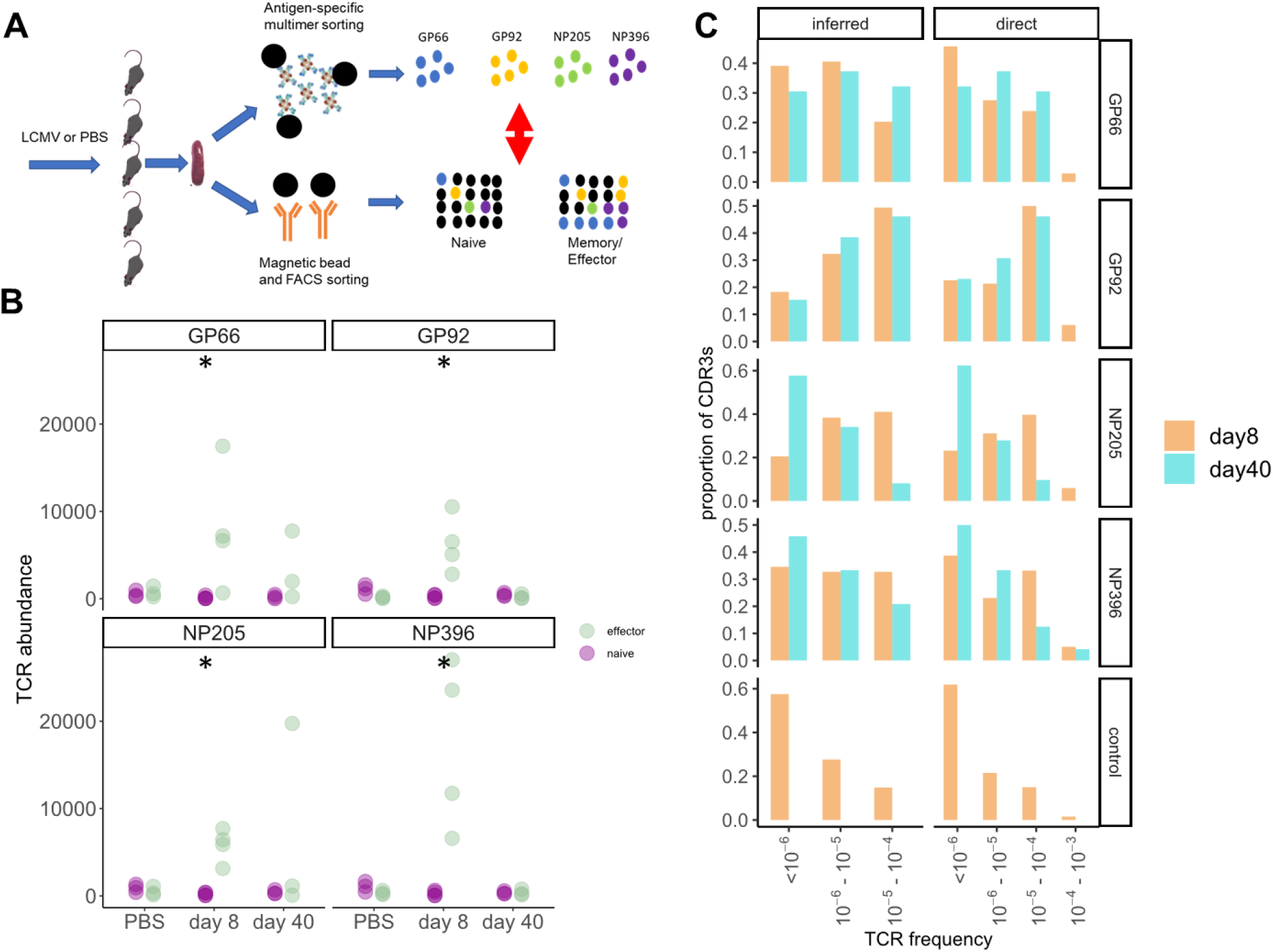
Abundance and frequency of LCMV specific TCR in naïve repertoires. A. A schematic of the experimental design. Mice were injected with PBS or LCMV and spleen T cells harvested at day 8 and day 40 post-infection. Epitope-specific T cells were isolated using specific tetramer sorting, and different subpopulations of bulk T cells were fractionated by a combination of FACS sorting and magnetic bead/antibody sorting. Both sets of T cells were processed for TCR sequencing and the overlap between the populations analysed. B. Total abundance of epitope-specific TCRs in the naïve (purple) or effector (green) TCR repertoires in individual mice. The abundance of all four epitopes was larger in the effector population in the day 8 infected group, than in the PBS group (* p < 0.001, Mann Whitney). All other pairwise comparisons were not significant. C Distribution histogram of frequency of epitope specific and control TCRs, estimated using either the statistical Poisson framework as in Fig 3 (left panels), or directly from TCR abundance data (right panels). Epitope-specific CDRs are split into CDRs that come up early (day 8, orange) and CDRs that come up later (day 40, cyan). The left column includes all epitope TCR which were not found within the bulk naïve population, and were assigned a frequency of less than 10^-6^.

There was a strong enhancement in the abundance of epitope specific TCRs in the effector compartment of LCMV infected mice, especially at day 8 post infection, compared to saline injected mice (Fig 5B). In contrast, although many epitope specific TCRs were detected in the naïve repertories, their abundance did not change between infected and PBS treated mice. Thus detection of epitope specific TCRs in the naïve repertories is unlikely to arise from contamination during the fractionation step.

We then examined the frequency distribution of the epitope specific TCRs in the naïve repertoires (Fig 5C). To evaluate the difference between TCRs that expand early (which might have higher precursor frequency) and TCRs which expand later, we classified the TCRs as day 8 (already detected in the bulk sequencing at day 8) or day 40 (not yet detected in bulk sequencing at day 8, but detected at day 40, Fig S14A). As shown in Fig 5C, in 3 out of 4 epitopes the early-rising antigen specific TCRs were enriched for high-abundance CDR3s, compared to the antigen-specific CDRs which arose at a later timepoint, or to a random sample of TCRs from the non-antigen specific repertoires (compare top four rows to bottom row). Because we used a quantitative TCR sequencing protocol, we were able to estimate frequency by two orthogonal methods. In the left column, we show the frequencies estimated using the Poisson sampling approach which we developed for the SARS-COV-2 data above. In brief, we determined the proportion of naïve repertoires in which we detected each TCR, and then used equation 2 to infer its frequency in the repertoire. Because we only had data from 10 mice, the granularity of this approach in this case was limited. In a second approach (right panels), we measured the average frequency of each TCR directly (counts/total number of TCR in sample), and then averaged the frequencies over all 10 mice. TCRs not detected were assigned a frequency less than 10^-6^ in both methods. Both methods of frequency estimation gave very similar results (Fig S14B), providing additional confidence in the statistical approach used for SARS-COV-2. Overall, these results suggest that high frequency TCRs specific for LCMV exist in the naïve repertoire prior to LCMV exposure.

## Discussion

The high prevalence and high degree of synchrony of the first wave of the SARS-COV-2 pandemic in the UK offered a rare opportunity to study the early dynamics of the T cell receptor repertoire following exposure to a natural novel viral infection. We identified a small number of TCRs which showed statistically significant changes in abundance within the first five weeks of our study, reasoning that viral exposure would lead to T cell clonal expansion and subsequent contraction. The set of TCRs identified in this way created a rapid transient wave of expansion, which peaked at or shortly after the first PCR+ SARS-COV-2 test. The number of expanded TCRs varied between individuals within a range of 18 to 912, and constituted only a very small fraction (1-2%) of the repertoire, consistent with the small proportion of activated T cells observed in blood using epitope specific tetramers or peptide specific T cell cytokine production (*48*, *49*).

The identification of the responding TCRs was unbiased and based on dynamics alone. The wave of expansion was not observed in PCR- controls, consistent with a SARS-COV-2 specific response. However, the rapid expansion of TCRs might arise from cytokine-driven antigen non-specific as well as antigen specific clonal expansion. Access to a number of large sets of CDR3s annotated as SARS-COV-2 specific, as well as similar sets annotated for CMV and EBV specificity by functional assays allowed us to interrogate the expanding set for antigen specificity. The expanded set of CDR3s were enriched for SARS-COV-2 specific annotated TCRs, although we cannot rule out that some by-stander activation of other memory cells may occur. The annotated CDR3s observed in our data set shared V gene usage with the V gene of the corresponding annotated TCRs. Furthermore, most expanded annotated TCRs were found in individuals expressing the same HLA allele as the reported restriction element of the annotated TCR. Both these observations build confidence in the functional annotation of the expanded TCRs as specific for SARS-COV-2.

The appearance of high frequency (> 100 TCRalpha/beta per million) T cells in the blood at the time of the first positive SARS-COV-2 PCR, or in some cases even before PCR+, needs to be considered in the context of the T cell precursor frequency prior to infection, and the known dynamics of T cell proliferation. The length of time between initial exposure and detectable SARS-COV-2 viremia (incubation period) has been estimated to be between 4 and 6 days (*50*–*53*). There is a lag between exposure and the arrival of antigen in the draining lymph node, and a further lag between TCR binding and the first cycle of T cell proliferation. Finally the time taken for a human T cell to complete the cell cycle is in the order of 12 - 14 hours (Phil Hodgkin, personal communication). There is therefore time for no more than a maximum of 10 T cell divisions which, without taking into account cell death, allows each cell to proliferate 2^10 (approximately 1,000-fold increase). The median frequency of the expanded TCRalpha and beta receptors prior to expansion must therefore be at least in the order of 1 in 10^7^. In a total pool of 10^11^ naïve T cells , this corresponds to a large (10,000) precursor T cell clone size.

In support of this hypothesis we were able to detect many of the expanded CDR3 sequences of the expanded TCRs in a significant proportion of pre-pandemic repertoires. We reasoned that TCRs which are present in many individuals (the public repertoire) must be present at high frequency in order to be observed in the small sample of blood which is typically sequenced. For example, in the Emerson data set of 786 repertoires collected and sequenced several years before the SARS-COV-2 pandemic, an average 100,000 TCR sequences are sequenced for each individual, out of an estimated total number of 10^11^ -10^12^ (*54*). We were therefore able to estimate the underlying frequency distribution of the 2648 expanded CDR3 TCRbeta sequences which we observed in two or more of the Emerson repertoires. As we predicted, the observed frequency distribution of these TCRs was between 10^-7^ and 10^-5^, with an average frequency of 1.8 * 10^-6^. At least 50% of the TCRs which expanded therefore had high precursor frequency in pre-COVID repertoires of healthy individuals.

We considered two possible hypotheses to explain the presence of high precursor frequency SARS-COV-2 specific T cells in the pre-pandemic repertoire. The first hypothesis is that these TCRs are present on cross-reactive memory T cells specific for homologous epitopes on circulating human coronaviruses. Similar cross-reactive T cells, especially directed at the conserved polymerase, have recently been described and may explain “abortive” infections with SARS-COV-2(*9*). However, an analysis of the epitopes recognized by the annotated early expanding TCRs identified in this study (which are predominantly directed at structural proteins) showed little conservation between SARS-COV-2 and the main circulating human coronaviruses (Fig 4). Therefore, while the existence of cross-reactive T cells is not in dispute, these cannot explain the early wave of SARS-COV-2 specific expansion documented in this study. In contrast, the epitopes recognized by the early expanding T cells were highly conserved (in most cases identical) between all different strains of SARS-COV-2. There is no evidence, therefore, that evolution has driven escape mutations within this set of T cell epitopes, similar to that seen for antibody. An important limitation of our study, however, is that the range of disease severity was very limited, and largely restricted to mild or asymptomatic disease. The potential role of early expanding T cells in limiting viral growth and hence pathology (*55*) cannot be investigated in this study, since all the infected HCW only developed mild disease .

The second hypothesis we explore is that the early SARS-COV-2-specific T cell response we observe can be attributed to TCRs present at high precursor frequency in the naïve compartment. Recent estimates suggest that the naïve repertoire may comprise 10^8^- 10^9^ different TCRs (*56*), but the frequency distribution is very broad with some TCRs present as high as 1 in 10^4^ (*41*). Several factors may drive differential frequency of different T cell receptors in the circulation, although the mechanisms remain incompletely understood. We could not measure the frequency of individual TCRs in the pre-infection naïve repertoires of the HCW from this cohort since we did not have sufficient stored PBMC to sort and sequence the naïve compartment prior to infection.

We reasoned that the phenomenon of rapidly responding high frequency T cells in the naïve repertoire might be a more general feature of the T cell adaptive immune system. We therefore investigated one of the classical most well studied models of virus infection, LCMV (*57*). We isolated epitope specific T cells expanding very early after infection (day 8) and searched for TCRs with identical CDR3s in the naïve, as well as effector repertoires of both immunized and unimmunized mice. The results we obtained were very similar to those we had observed with SARS-COV-2, but in this case we were able to directly test our principal hypothesis that T cells with high endogenous frequency are present within the naïve compartment, and contribute significantly to the early wave of T cell responses following infection. Moreover, we could detect a difference in the distribution of frequency in the naïve repertoire between cells that arise early and cells that are detected later in the infection, further supporting our hypothesis that early-expanding cells have higher precursor frequencies in the naïve compartment.

The design of the COVIDsortium study imposed some limitations. The protocol of the study, which was set up as rapidly as possible to capture the first wave of the SARS-COV-2 pandemic in the UK, did not incorporate collection of sufficient number of viable cells at the early time points to carry out functional antigen-specific assays. Although we did not observe a wave of early TCR expansion/contraction in PCR- seronegative individuals, we cannot therefore exclude that some of the expanding TCRs were bystander memory cells to other antigens triggered by cytokine production. Furthermore, as discussed above, the same limitations on numbers of stored cells did not allow us to test directly our principal hypothesis that the early expanding TCRs arose from high frequency pre-exposure naïve T cell clones.

Despite these limitations, our study raises the intriguing possibility that the human immune system contains a population of high frequency precursor T cells which can provide a very rapid response to viral infections. These rapid responses do not require cross-reaction with circulating human coronaviruses, but are associated with pre-existing high frequency precursors. Further studies will be required to determine whether these are high frequency naïve T cells, or represent random cross-reactivity of the memory pool. The ability to stimulate this rapid response may be an important factor in designing future vaccines to limit the growth and hence pathogenesis of respiratory viral infections.

## Acknowledgments

Funding for COVIDsortium was donated by individuals, charitable Trusts, and corporations, including Goldman Sachs, Kenneth C. Griffin, The Guy Foundation, GW Pharmaceuticals, Kusuma Trust, and Jagclif Charitable Trust, and enabled by Barts Charity with support from UCLH Charity. R.K.G. is funded by the National Institute for Health Research (DRF-2018-11-ST2-004). J.C.M., C.M., and T.A.T. are directly and indirectly supported by the University College London Hospitals NHS Foundation Trust (UCLH) and Barts NIHR Biomedical Research Centers and through the British Heart Foundation (BHF) Accelerator Award (AA/18/6/34223). MM is supported by a Cancer Research UK studentship. T.A.T. is funded by a BHF Intermediate Research Fellowship (FS/19/35/34374). M.N. is supported by the Wellcome Trust (207511/Z/17/Z) and by NIHR Biomedical Research Funding to UCL and UCLH. M.K.M. is supported by UKRI/NIHR UK-CIC and Wellcome Trust (214191/Z/18/Z). G.S.T. is supported by a UK Medical Research Council Clinician Scientist Fellowship (MR/N007727/1). R.J.B. and D.M.A. are supported by UKRI/MRC Newton (MR/S019553/1, MR/R02622X/1, MR/V036939/1, MR/W020610/1), NIHR Imperial Biomedical Research Center (BRC): ITMAT, Cystic Fibrosis Trust SRC (2019SRC015), NIHR EME Fast Track (NIHR134607), NIHR Long Covid (COV-LT2-0027), Innovate Uk (SBRI 10008614), and Horizon 2020 Marie Skłodowska-Curie Innovative Training Network (ITN) European Training Network (no. 860325). A.M. is supported by Rosetrees Trust, The John Black Charitable Foundation, and Medical College of St Bartholomew’s Hospital Trust. B.M.C. is supported by the Rosetrees Trust, the NIHR UCLH BRC, and the Wellcome Trust. Support was also provided by UK Medical Research Council (T.D, Y.P), Chinese Academy of Medical Sciences (CAMS) Innovation Fund for Medical Sciences (CIFMS), China (grant number: 2018- I2M-2-002) (T.D, Y.P, X.Y, G.L, S.L.F ), National Institutes of Health, National Key R&D Program of China (2020YFE0202400) (T.D), NIHR Oxford Biomedical Research Centre (J.C.K, A.J.M)

## Author contributions

Conceptualization: MN, BC
Data acquisition and analysis: MM, YP, CT, MM , GN, SB , AC, JR, MW, XY, GL, SLF, TR
Visualization: CT, MM, TP
Funding acquisition: JCM, MN
Project administration: CM, TT, JCM
Supervision: TD, FB, AMcK,CM, TT, MN, BC, EG, SR-Z, CP, JG, AS, TB, DMA, RJB, MKM
Writing – original draft: MM, MN, BC
Writing – review & editing: CP, AS, RB, EG, FB, CT, BC, MM, MN, MKM, AM

## Competing interests

Authors declare that they have no competing interests.

## Data and materials availability

The COVID_sortium TCR sequencing data are available at NCBI Short Read Archive accession number SUB9362448 (https://www.ncbi.nlm.nih.gov/sra).). The mouse TCR sequences have been submitted to the Sequence Read Archive under identifier PRJNA771880. https://www.ncbi.nlm.nih.gov/Traces/study/?acc=PRJNA771880&o=acc_s%3Aa All other data are available in the main text or the supplementary materials. All code for analysis and generating individual figure panels are available from BC on request.

## Notes

### Competing Interest Statement

The authors have declared no competing interest.

