## Supplementary material for "Large clones of pre-existing T cells drive early immunity against SARS-COV-2 and LCMV infection": Methods and supplementary

Supplementary Materials

Materials and Methods

**Ethical approval.** The COVIDsortium study was approved by a UK Research Ethics Committee (South Central - Oxford A Research Ethics Committee, reference 20/SC/0149). All participants provided written informed consent. The animal studies were performed according to regulations formulated by The Weizmann Institute’s Animal Care and Use Committee.

**COVIDsortium study design.** We undertook a case control study nested within our COVIDsortium health care worker cohort. Participant screening, study design, sample collection, and sample processing have been described in detail previously (*3*, *17*) and the study is registered at ClinicalTrials.gov (NCT04318314). Briefly, healthcare workers were recruited at St Bartholomew’s Hospital, London, UK in the first week of lockdown in the United Kingdom (between 23^rd^ and 31^st^ March 2020). In London, the case-doubling time in March, 2020 was approximately 3–4 days. The number of nasal swabs testing positive for SARS-CoV-2 in our study peaked at March 23rd to 31st, 2020 suggesting that infections peaked on or around March 23rd, 2020, the day of UK lockdown. We thus observed approximately synchronous infections coincident with the peak epidemic transmission in London at the start of the study, UK lockdown on March 23rd and therefore used this as the benchmark starting point for our analysis in the first wave. Participants underwent weekly evaluation using a questionnaire and biological sample collection (including serological assays) for up to 16 weeks when fit to attend work at each visit, with further follow up samples collected at 6 months.

Participants with available blood RNA samples who had PCR-confirmed SARS-CoV-2 infection (Roche cobas® diagnostic test platform) at any time point were included as ‘cases’. Six participants without evidence of SARS-CoV-2 infection on nasopharyngeal swabs and who remained seronegative by both Euroimmun anti-S1 spike protein and Roche anti-nucleocapsid protein throughout follow-up were included as uninfected controls.

**Genotyping**. All samples were HLA typed as described in detail previously (*18*).

**T cell receptor sequencing and analysis** The α and β chains of the TCR repertoire were sequenced from all time points for which RNA was available within the first 4 weeks of the study for all participants who were PCR+ at any time point, and for six randomly selected individuals who remained PCR- and seronegative throughout the study. The pipeline introduces unique molecular identifiers attached to individual cDNA molecules using single-stranded DNA ligation The UMI allows correction for sequencing error PCR bias, and provides a quantitative and reproducible method of library preparation. Full details for both the experimental TCRseq library preparation and the subsequent TCR annotation (V, J and CDR3 annotation) using Decombinator V4 are published in (*19*, *20*). The Decombinator software is freely available at https://github.com/innate2adaptive/Decombinator

Expanded TCRs were defined as any TCR which changed significantly between any two time points. The significance boundaries (shown as blue dotted lines in Figure 1A) were defined as the maximum TCR abundance which might be observed at time 2, given its abundance at time 1, given Poisson distribution of counts with p < 0.0001, to give a False Discovery Rate of <1 in 1000. TCR abundances are normalised for total number of TCRs sequenced in each sample, and expressed as counts/million. MAIT TCRs were defined as any TCR alpha containing TRAV1-2 paired with TRAJ12, TRAJ20 or TRAJ33. iNKT TCRs were defined as TCRs containing TRAV10 paired with TRAJ18.

**TCR clustering**. The pairwise distance between expanded or control CDR3 sequences was measured by measuring the number of shared triplet amino acid motifs between the two CDR3s, and normalizing by the length of the CDR3 to give a similarity metric between 1 (identity) and zero (no sharing). The analysis was done using the stringdot and kernelMatrix functions in the kernlab package V0.9-31 (*21*). The similarity matrix was plotted as a graph (layout format fruchterman-reingold), using the package igraph version 1.2.8 (*22*) implemented in R. Each CDR3 is represented as a node, and nodes are connected by an edge if the triplet similarity is greater than a given threshold. The thresholds used for clustering were 0.72 for CDR3α and 0.76 for CDR3β, to give a FDR in similar size control sets of 1 in 10,000.

**TCR annotation.** TCR sequences of SARS-CoV-2 specific T cells were obtained as previously described (*23*). In brief, antigen-specific CD4+ and CD8+T cells were stained with peptide-MHC Class I pentamer and peptide-MHC class II tetramer, respectively, and were then sorted by flow cytometry for single cell RNASeq. T cell receptor sequences of each single cell were reconstructed from Smartseq2 scRNA-seq FASTQ files using MiXCR v.3.0.13 (*24*), or extracted from 10x VDJ sequencing using cellranger vdj (Cellranger). Deep sequencing of TCR repertories of T cell clones was carried out using a SMARTer Human TCR a/b Profiling Kit (Takara) following the supplier’s instructions.

In order to assess the sensitivity of the SARS-CoV-2 specific T cells, SARS-CoV-2 specific T cell clones were generated as previously (*25*). T cell clones were then co-cultured with target cells loaded with peptide at titrated concentration. Cytokine production of each T cell clone was then assessed by IFN-gamma ELISpot or intracellular cytokine staining. EC50 of the T cells were then calculated by Prism using nonlinear regression with variable slope. T cell clones with lower EC50 are more sensitive to antigen stimulation.

**Homology mapping of SARS-COV-2 epitopes**. We used blastp from BLAST+ v2.11.0 to compute the sequence homology of the 59 unique epitope sequences against a database of protein sequences derived from all seven species of existing human-associated coronaviruses (HCoVs). We used the same parameters as our previous study, which are optimised for short query sequences (*26*). Briefly, -task, -qcov_hsp_perc, -num_alignments, -evalue were set to blastp-short, 99, 109 and 2 × 109, respectively.

To construct the blastp database, we retrieved all genome records for HCoV-NL63 (taxid:277944; n = 71), HCoV-229E (taxid:11137; n = 41), HCoV-HKU1 (taxid:290028; n = 33) and HCoV-OC43 (taxid:31631; n = 206) on NCBI Virus. Only accessions for genomes isolated from human hosts and flagged as ‘complete’ were retained. Protein sequences for all accessions were downloaded using the online Batch Entrez utility (https://www.ncbi.nlm.nih.gov/sites/batchentrez). Separately, we retrieved the WIV01 reference and a random sample of 50 genomes for each existing variant of concern (VoC): Alpha, Beta, Gamma, Delta, and Omicron from the 15th June Audacity release on GISAID (*27*, *28*). Protein annotations of WIV01 were obtained from GenBank (accession: NC_045512.2) and used to obtain protein sequence annotations from the SARS-CoV-2 genomes. To do so, we aligned the SARS-CoV-2 genomes against WIV01 using the Augur v14.0.04 wrapper for MAFFT v7.490 (*29*, *30*). Protein sequences were translated from the aligned SARS-CoV-2 genomes using the trans function as part of the Ape v5.6.2 package in R (*31*). All custom scripts used to perform these analyses are hosted on GitHub (https://github.com/cednotsed/early_tcell_x-reactivity.git). All public databases were accessed on 20th June 2022.

**Emerson data set.** The Emerson dataset was downloaded from the link provided in (*32*). We used all 786 samples (cohort and control), and computed for each CDR3 its sharing level in the cohort. The sharing level of an amino acid CDR3 sequence is defined by the number of repertoires in the cohort that contain the given CDR3.

**LCMV infections.** Seven 5 weeks old (Envigo) C57BL/6 female mice were injected intraperitoneally with 2 x 10^5^ units plaque forming units of the Armstrong LCMV strain. Mice were sacrificed after 8 or 40 days of infection, and spleens were collected. An additional 3 mice were injected with saline and sacrificed on day 8. All mice were maintained in specific pathogen-free conditions.

**Mouse T cell isolation**. Spleens were dissociated with a syringe plunger and single cell suspensions treated with ammonium-chloride potassium lysis buffer to remove erythrocytes. Bone marrows were extracted from the femur and tibiae of the mice and washed with PBS. Samples were loaded on MACS column (Miltenyi Biotec) and T cells were isolated according to manufacturer’s protocol. Bone marrows cells were purified with CD3+ T isolated kit (CD3ε MicroBead Kit, mouse, 130-094-973, Miltenyi Biotec). Splenic CD4+ and CD8+ cells were purified in two steps: (1) CD4+ positive selection (CD4+ T Cell Isolation Kit, mouse, 130-104-454, Miltenyi) (2) the negative cell fractions were further selected for the CD8+ positive cells (CD8a+ T Cell Isolation Kit, mouse, 130-104-07, Miltenyi Biotec). For the tetramer binding reaction, we pooled splenocytes from previously infected mice (5 mice after 8 days of infection) and purified their T cells using the T cell isolation kit (Pan T Cell Isolation Kit II, mouse, 130-095-130, Miltenyi Biotec).

**Mouse Flow cytometry analysis and cell sorting** The following fluorochrome-labeled mouse antibodies were used according to the manufacturers’ protocols: PB or Percp/cy5.5 anti -CD4+, PB or PreCP/cy5.5 anti- CD8+, PE or PE/cy7 anti- CD3+, APC anti-CD62L, Fitc or PE/cy7 anti- CD44 (Biolegend). Cells were sorted on a SORP-FACS-AriaII and analysed using FACSDiva (BD Biosciences) and FlowJo (Tree Star) software (*33*). Sorted cells were centrifuged (450g for 10 minutes) prior to RNA extraction.

**LCMV-tetramer staining and FACS sorting** Three monomers (NIH Tetramer Core Facility) with different LCMV epitopes were used: MHCI- NP396-404(H-2Db), MHCI- NP205-212(H-2Kb), MHCI- GP92-101 (H-2Db) and MHCII-GP66–77(H-2Ab). Tetramers were constructed by binding biotinylated monomers with PE/APC – conjugated- streptavidin (according to the NIH protocol). Purified T cells were stained with FITC anti-CD4+ and PB anti-CD8+ followed by tetramer staining (two tetramers together), for 30 min at room temperature (0.6ug/ml). CD8+ or CD8+ epitope-specific cells were sorted from single-positive gates for one type of tetramer (Fig S1).

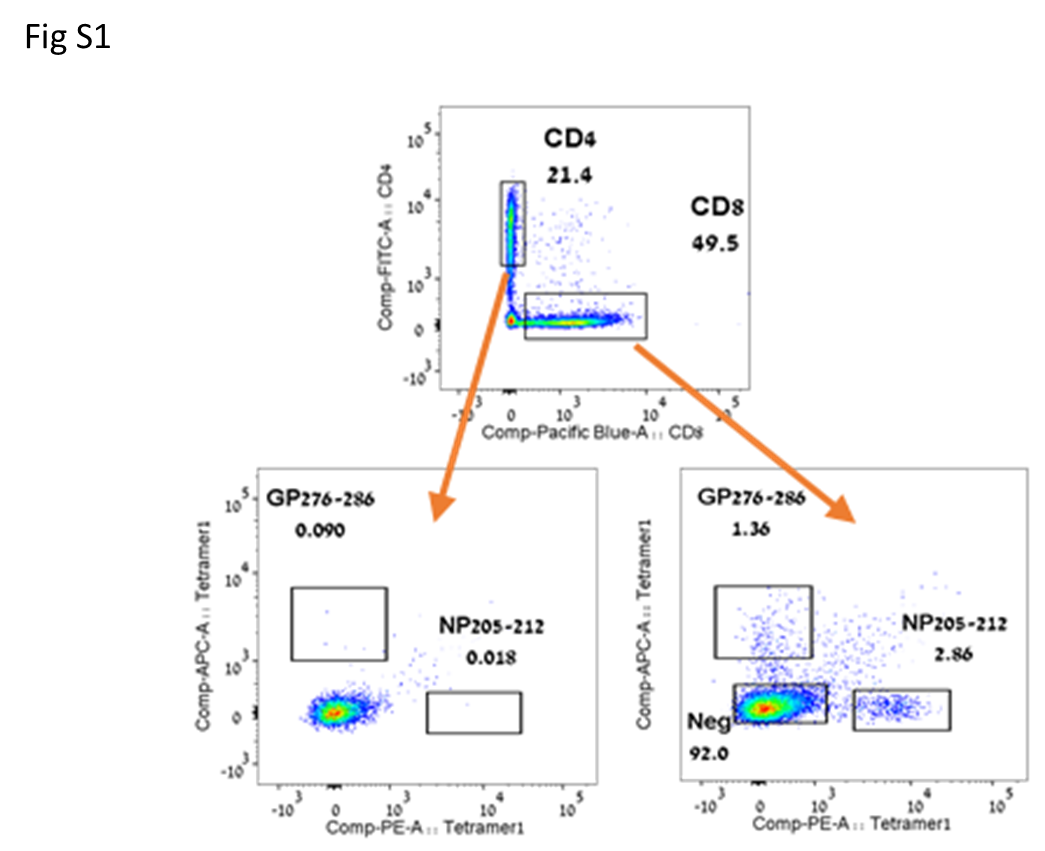

Fig S1 Representative sort plot for isolation of CD8+ LCMV tetramer positive cells.

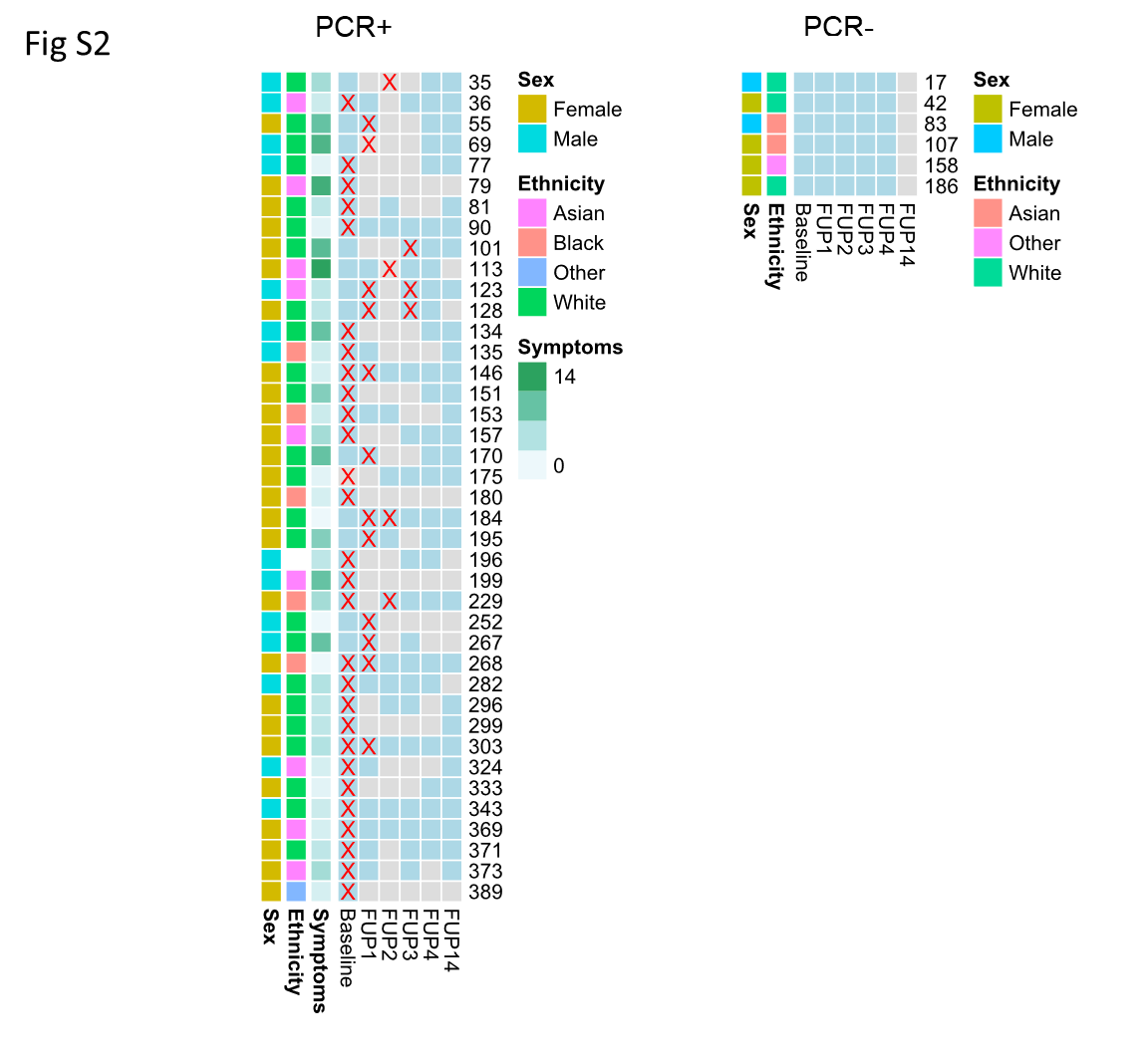

Fig S2 A. Summary of samples from baseline, and follow-up (FUP) from the COVIDsortium study. Each row represents one individual, and the anonymized study ID is shown to the right of the row. The samples for which RNA was available are shown in light blue. The red X shows the time of first PCR+ test. Basic demographics are shown for each individual.

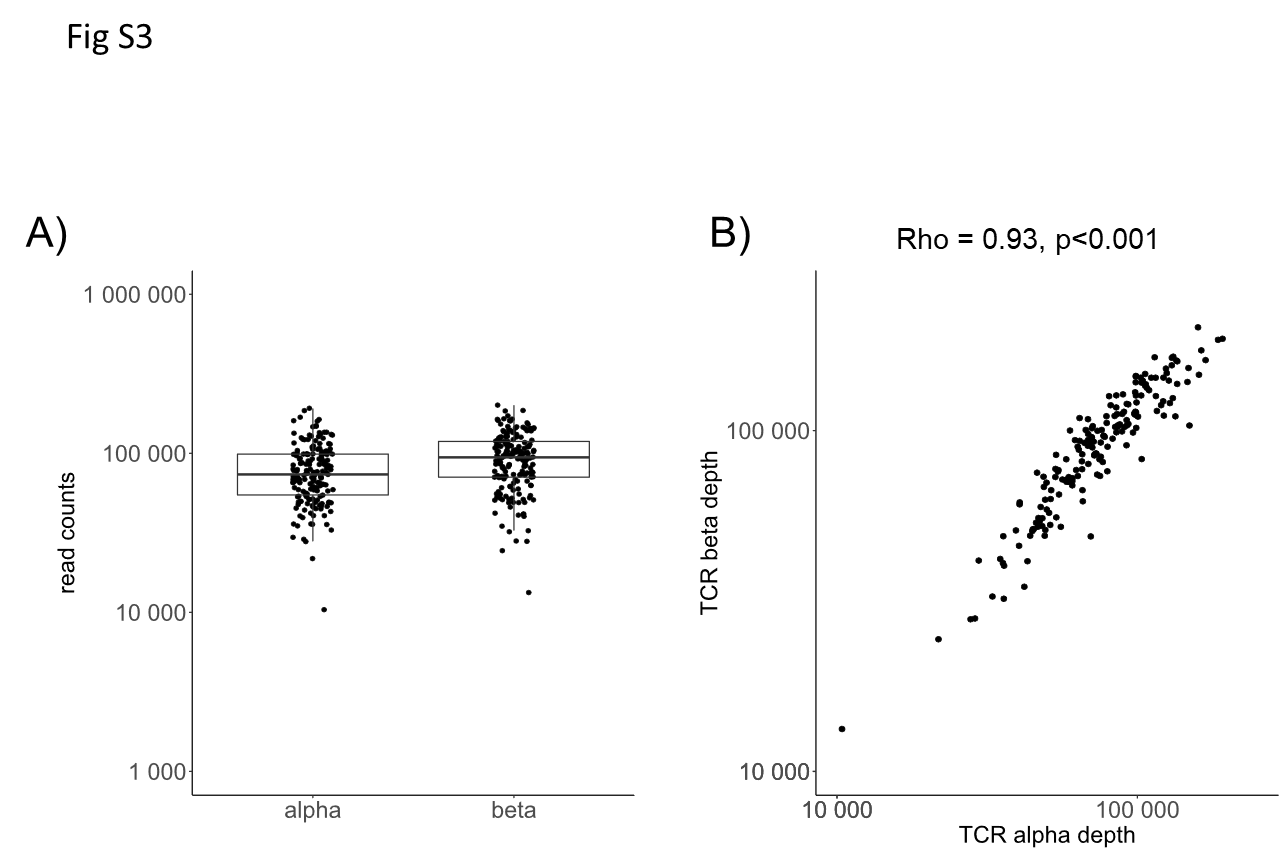

Fig S3 A) Total number of TCRs sequenced per sample. B) Total number of beta versus total number of alpha per sample.

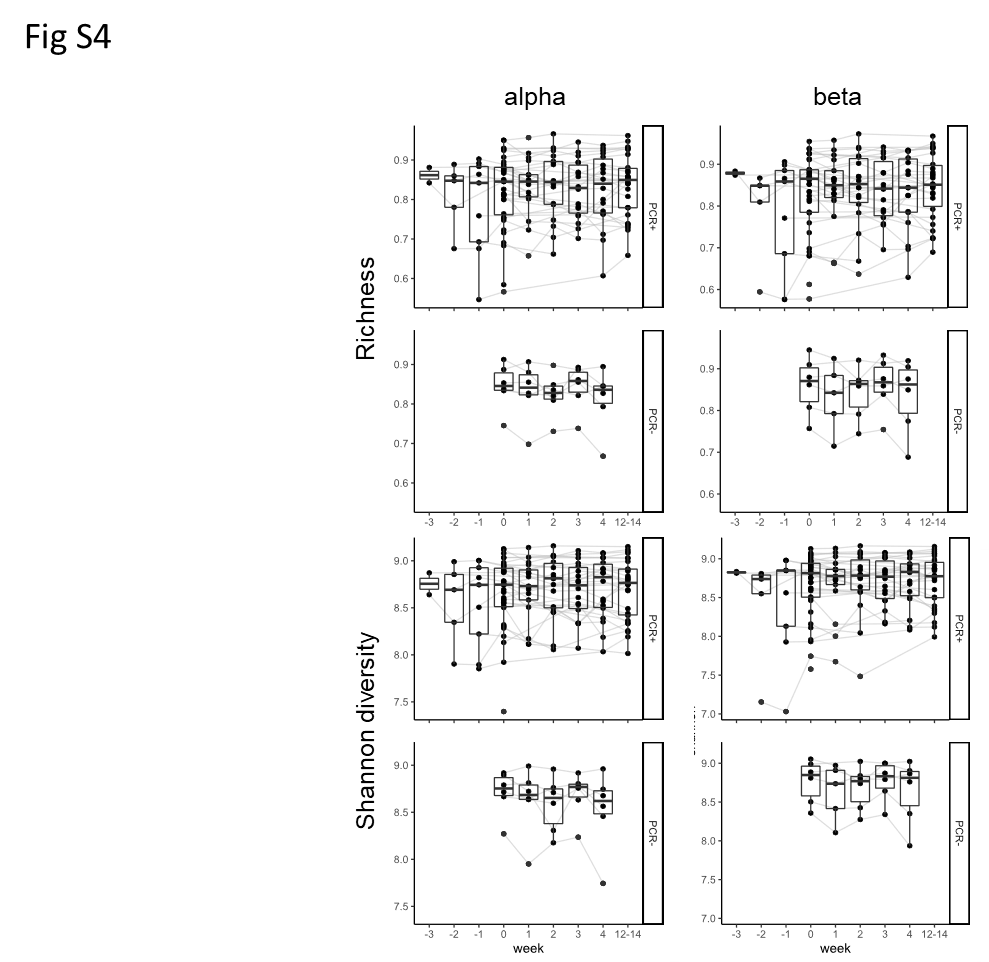

Fig S4. The richness and Shannon diversity of the TCR repertoires at each time point. For PCR+ individuals, the x-axis is rescaled relative to the week at which they first became PCR+ (this is week 0). For PCR- individuals the weeks correspond to baseline and subsequent follow-ups at weeks 1-4.

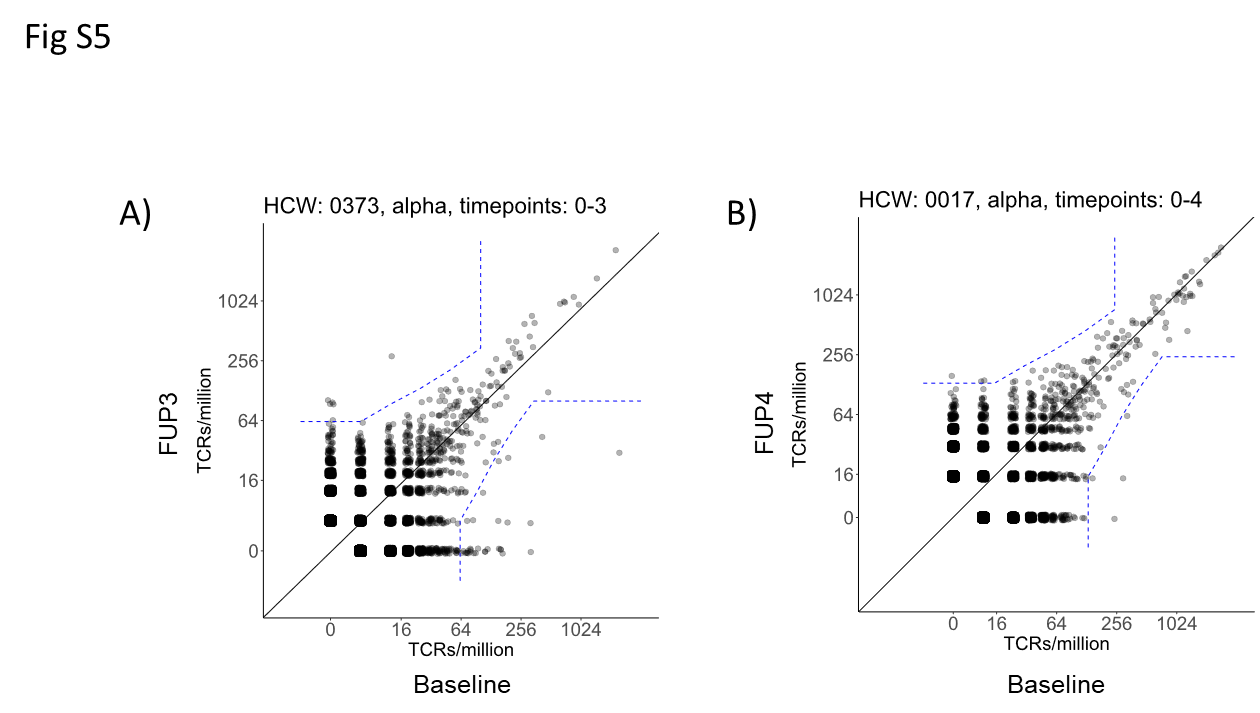

Fig S5

A). An example of a pairwise comparison showing TCRs contracting between baseline and FUP3 timepoints. The individual (ID 373) was PCR+ at baseline. Each point is an individual TCR sequence, and the plot shows abundance at FUP3 versus abundance at Baseline. All abundances are normalized to number of TCRs per million. The dashed blue line indicates the significance threshold calculated as described in M&M. All TCRs which fall outside the dashed line are considered as expanded (contracted). B) As in A, comparing repertoires at baseline and FUP4 in a control individual (ID 17) who did not become PCR+ or seroconvert during the study. Note the small number of expanding (contracting) TCRs.

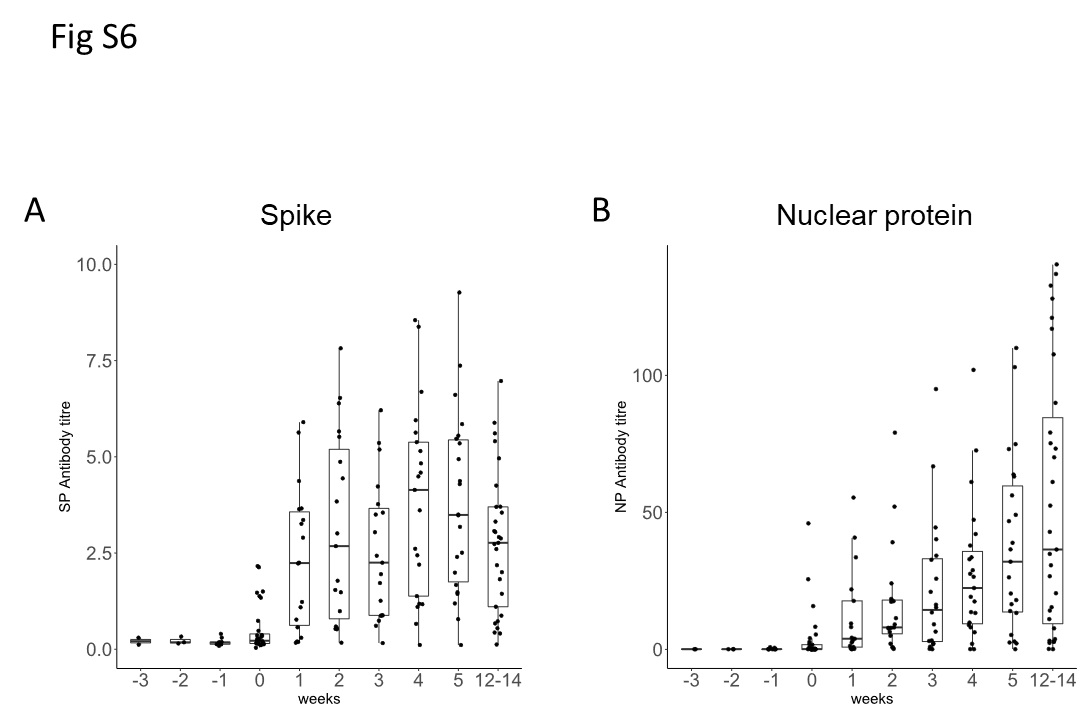

Fig S6. The kinetics of anti-SARS-COV-2 antibody in serum of the HCW who became PCR+. The y axis is an arbitrary scale of antibody titre.

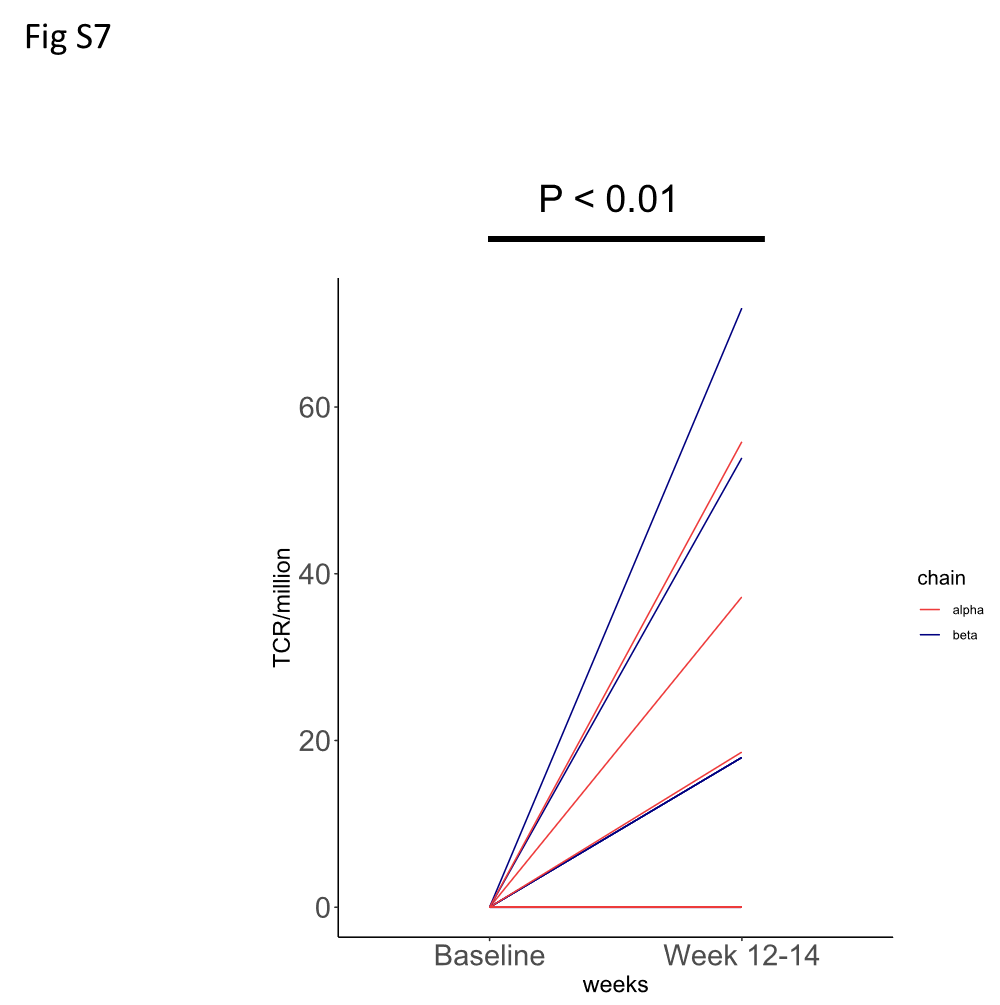

Fig S7 Comparison of TCR abundance at baseline (pre-PCR+, week -3) and week 14 for HCW 101. TCRalpha is shown in red, TCRbeta in blue.

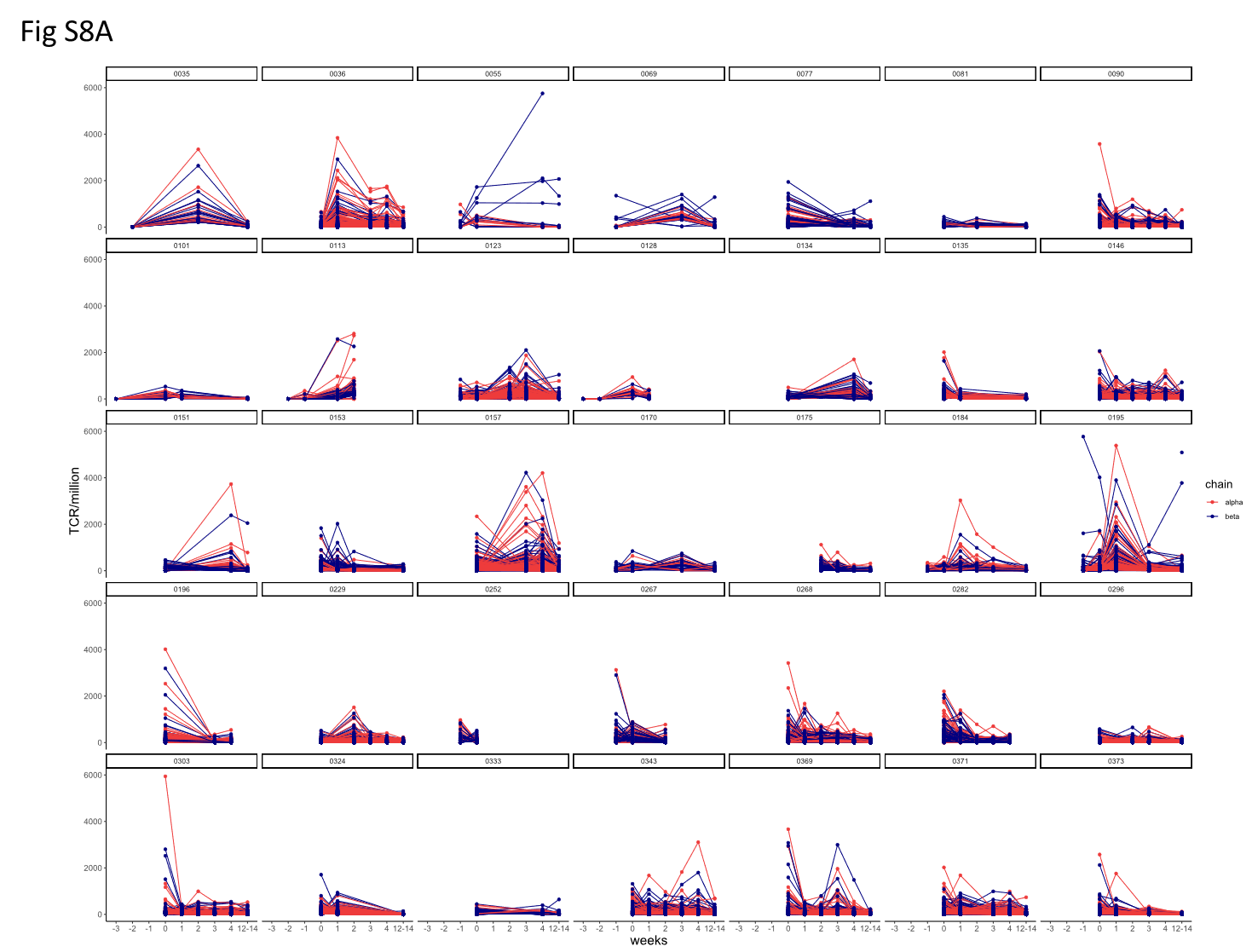

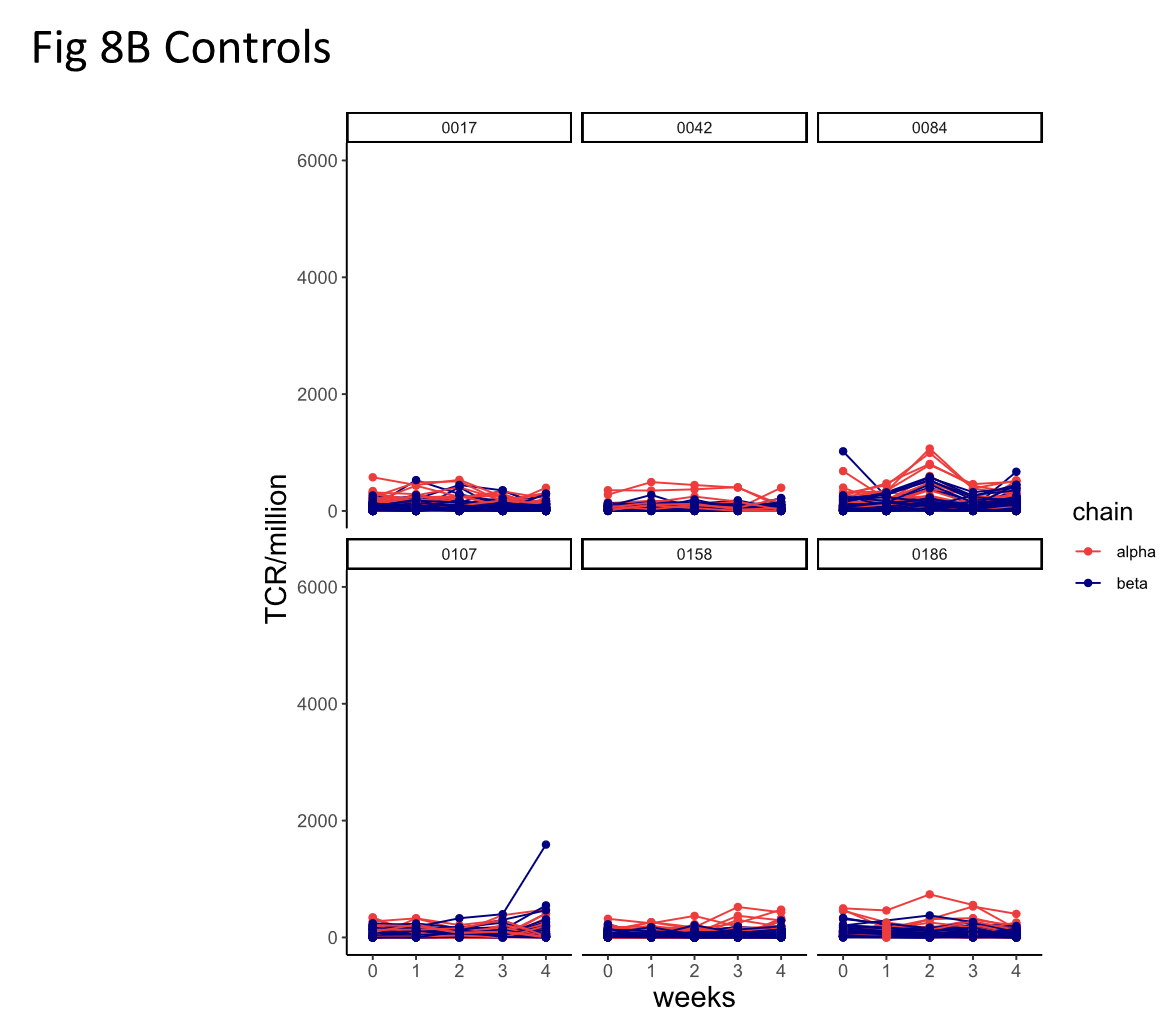

Fig S8

A. Kinetics of changing TCRs in individual repertoires of each PCR+ HCW.
B. same for controls (same scale for comparison)

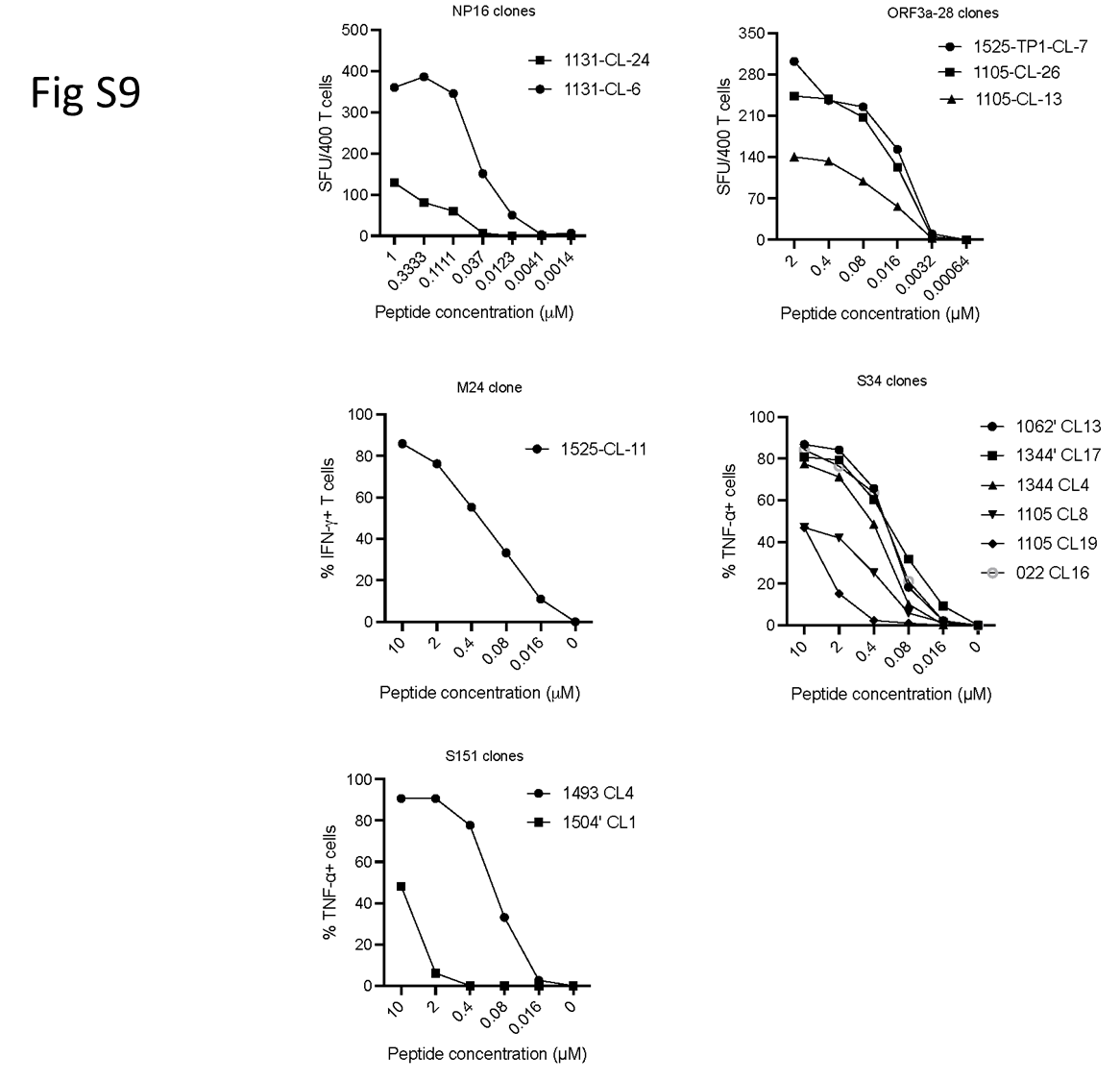

Fig S9 Evaluation of antigen sensitivity of SARS-CoV-2 specific T cells. SARS-CoV-2 specific T cell clones were generated as previously described <https://doi.org/10.3389/fimmu.2015.00287>. T cell clones were co-cultured with target cells loaded with peptide at titrated concentration and cytokine production of each T cell clone was assessed by IFN-γ ELISpot or intracellular cytokine staining.

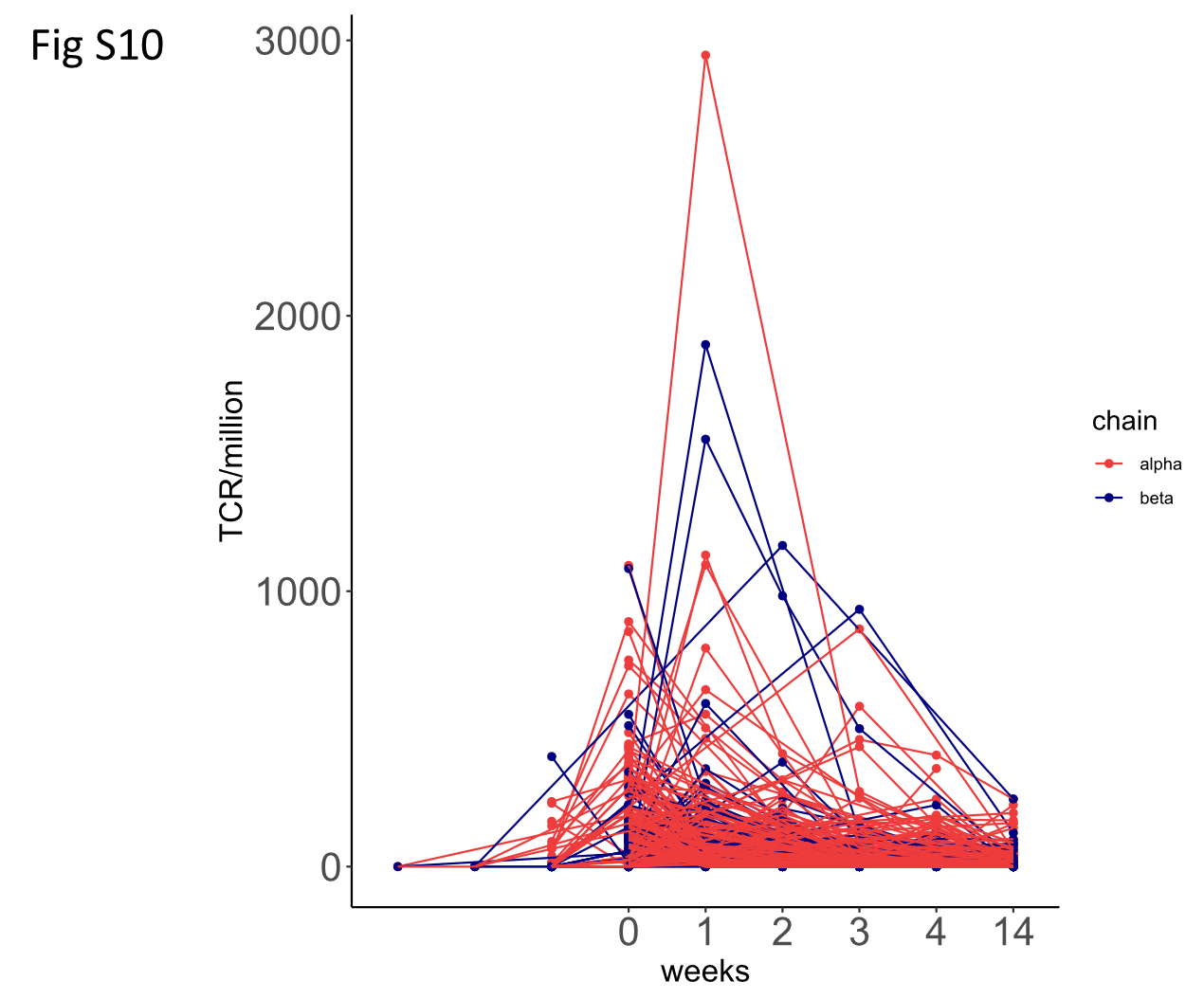

Fig S10. The time course or TCRs which are both expanded and annotated as recognising SARS-COV-2 antigens.

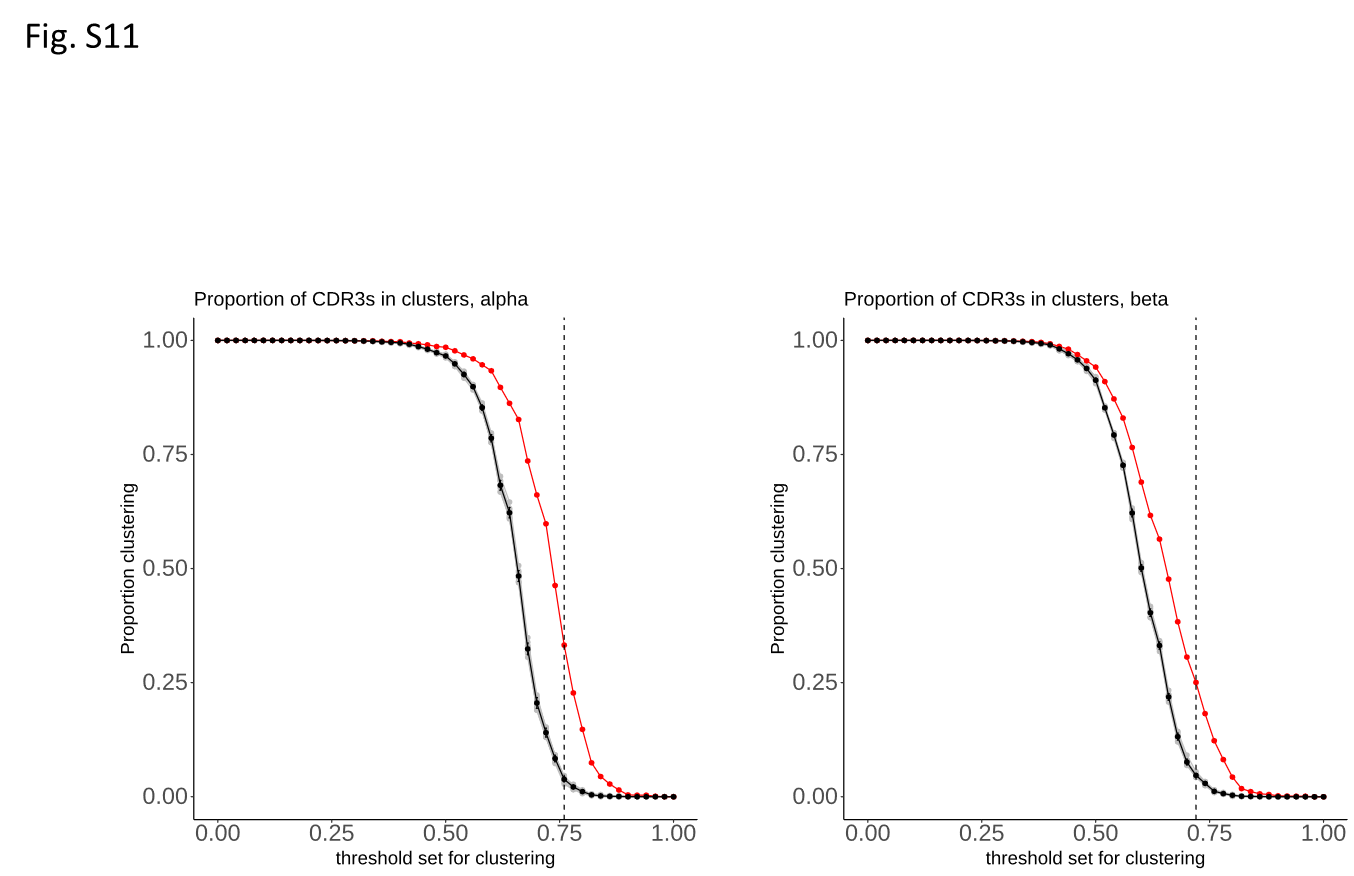

Fig S12. The proportion of TCRalpha and TCRbeta within a cluster (cluster size > 1) at different similarity thresholds. The red line shows the SARS-COV-2 expanded TCRs; the black line shows the mean of 10 sets of control (non-expanded) TCRs. The vertical dotted line shows the threshold used in the plots shown in Fig 2F.

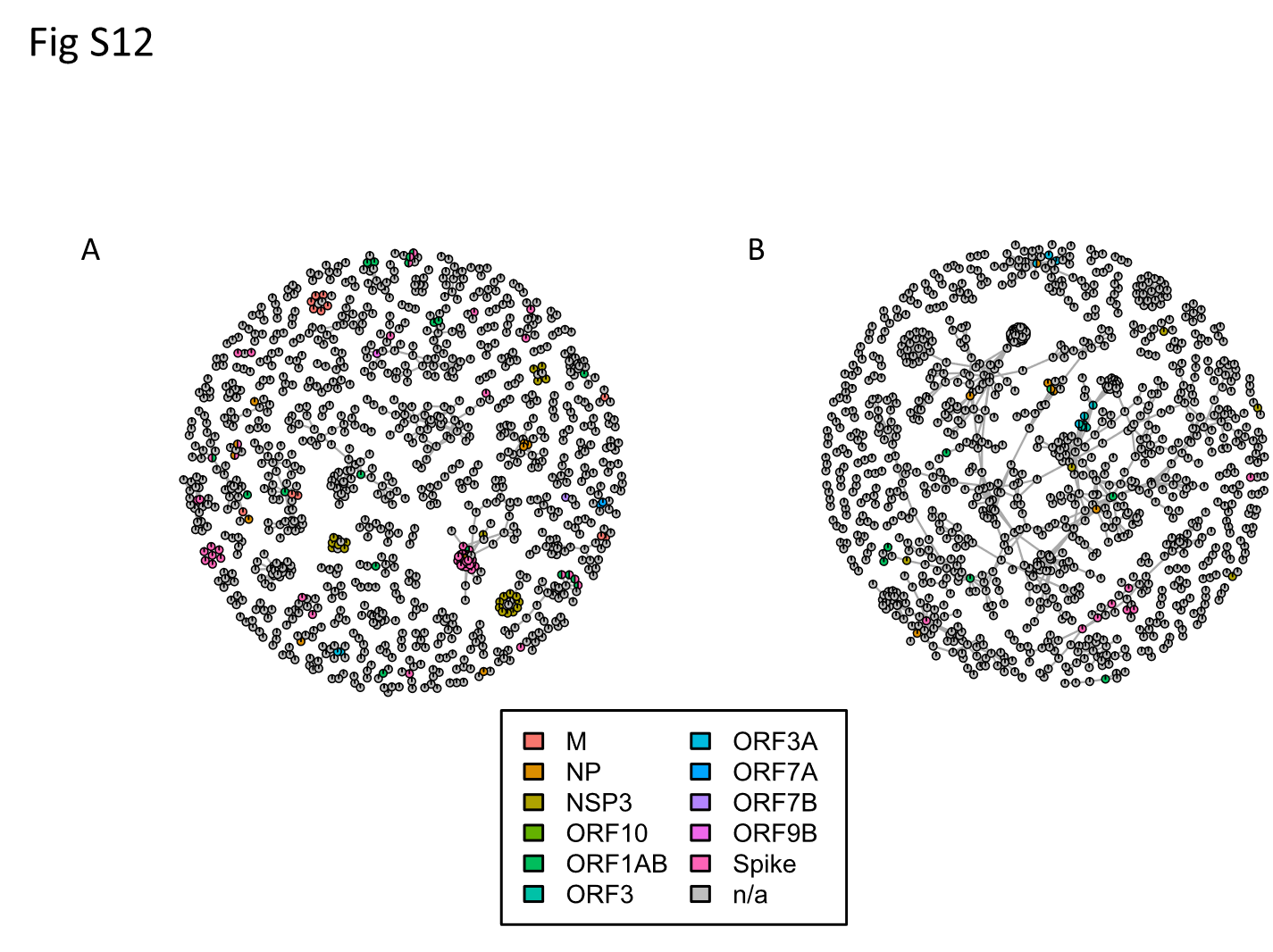

Fig S12 Clustered expanded TCRalpha (A) or TCRbeta (B) sequences (clusters with greater than 3 nodes are shown) coloured according to target antigen. The grey nodes represent unannotated TCRs for which the antigen is not known.

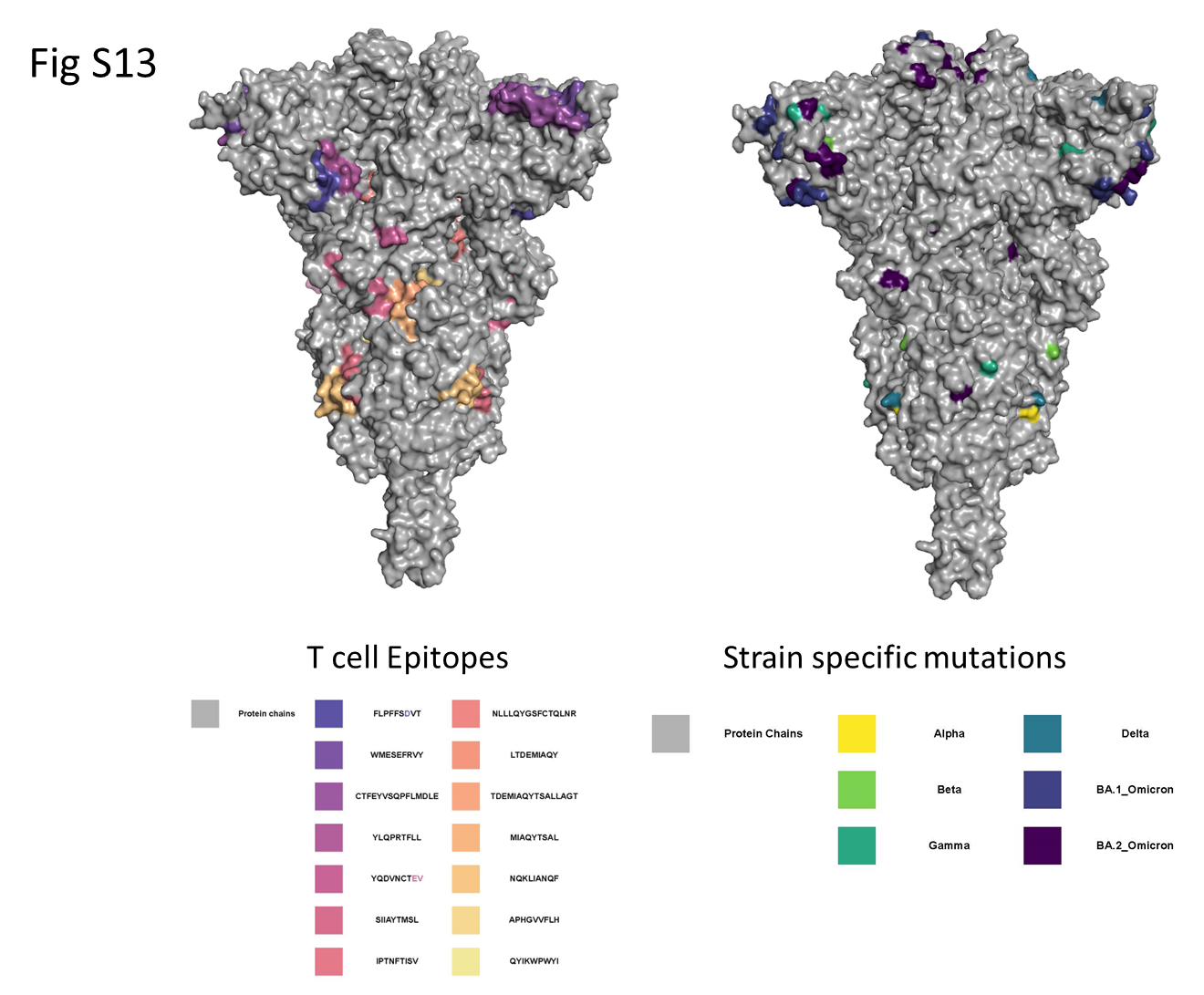

Fig S13 Location of defined spike T cell epitopes and strain-specific mutations within the spike protein structure (data from Fig 4). .

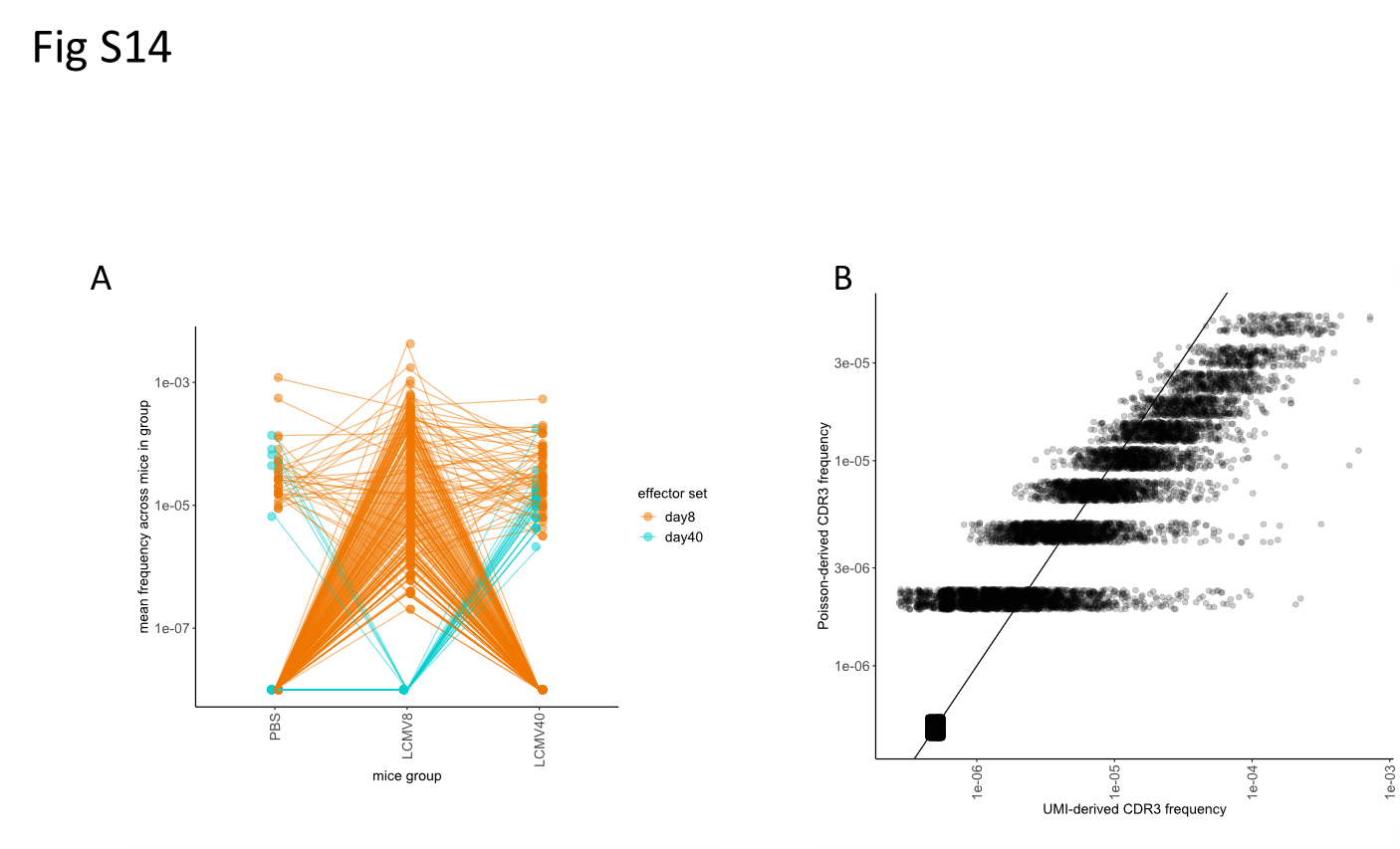

Fig S14

A. Definition of effectors as expanded day 8 (orange) or expanded day 40 (cyan). The average frequency of each CDR3 across the 3 (or 4) mice at each timepoint is shown.
B. The naïve frequency of each CDR3 inferred with a method based on Poisson statistics and calculated from a quantitative sequencing protocol correlate with each other (Pearson R = 0.78, p-value < 2.2e-16)

Complete list of COVIDsortium investigators.

COVIDsortium investigators (Surnames in alphabetical order)

Hakam Abbass

Aderonke Abiodun

Mashael Alfarih

Zoe Alldis

Daniel M Altmann

Mervyn Andiapen

Jessica Artico

Joao Augusto

Georgina L Baca

Anish Bhuva

Alex Boulter

Ruth Bowles

Rosemary J Boyton

Olivia Bracken

Timothy Brooks

Natalie Bullock

Gabriella Captur

Benny Chain

Nicola Champion

Carmen Chan

Jorge Couto de Sousa

Xose Couto-Parada

Marie-Teresa Cutino-Moguel

Rhodri H Davies

Keenan Dieobi-Anene

Karen Feehan

Malcolm Finlay

Marianna Fontana

Nasim Forooghi

Joseph M Gibbons

Derek Gilroy

Peter Griffiths

Rishi K Gupta

Matt Hamblin

Lauren M Hickling

Aroon D Hingorani

Lee Howes

Ivie Itua

Victor Jardim

Melanie Jensen

Meleri Jones

George Joy

Vikas Kapil

Jonathan Lambourne

WY Jason Lee

Mala K Maini

Vineela Mandadapu

Charlotte Manisty

Aine McKnight

Katia Menacho Medina

Celina Mfuko

Oliver Mitchelmore

James C Moon

Mahdad Noursadeghi

Ben O’Brien

Ben Ollivere

Corinna Pade

Susana Palma

Kush Patel

Ruth Parker

Brian Piniera

Alicja Rapala

Amy Richards

Mathew Robathan

Genine Sambile

Amanda Semper

Andreas Seraphim

Angelique Smit

Michelle Sugimoto

George D Thornton

Thomas A. Treibel

Arthur Tucker

Ana Valdes

Jessry Veerapen

Mohit Vijayakumar

Timothy Warner

Sophie Welch

Dylan Williams

Theresa Wodehouse

Lucinda Wynne

Dan Zahedi
